# Photosoftening Macroporous Hydrogels for Dynamic Tissue Engineering

**DOI:** 10.64898/2026.07.13.737088

**Authors:** Golnaz Navidi, Ben Canter, Ella Morris, Teresa L. Rapp

## Abstract

With the push towards accessible benchtop models to capture biological events, many researchers are reaching for hydrogel platforms for 3D tissue engineering ex vivo. Recapitulating the dynamic mechanical environment cells experience in vivo requires dynamic hydrogel scaffolds whose mechanical properties can be reprogrammed with spatiotemporal precision. Here we describe a chemically simple hydrogel platform that undergoes visible-light photosoftening via a ruthenium-based photocleavable crosslinker, leveraging tetrazine-norbornene inverse electron demand Diels Alder (iEDDA) click chemistry between RuTetrazine crosslinker and norbornene-modified hyaluronic acid (NorHA). Nitrogen gas evolved during this reaction is repurposed as an intrinsic porogen, nucleating macropores (55–175 µm) directly during gelation. Initial stiffness (1.5–10 kPa) and softening extent (from 50%-100% drop in storage modulus) are independently tunable through polymer and crosslinker composition. We have found RuTetrazine to be non-mutagenic and non-toxic (>80% live cell populations) once network-bound (IC50 = 0.27 mM). In a cell-instructive network co-crosslinked with an MMP-RGD-bearing peptide, human mesenchymal stromal cells (hMSCs) photosoftened in situ (2.27→0.54 kPa, ∼76%) spread approximately six-fold relative to stiff controls (∼6,500 vs. ∼1,100 µm^2^, p < 0.0001). This work demonstrates a synthetically accessible photocleavable crosslinker and a simple, macroporous hydrogel for modulating dynamic mechanical cues in three dimensions.

## 1. Introduction

Hydrogel scaffolds are used in tissue engineering to (1) replicate tissue-relevant mechanical properties, including stiffness and viscoelasticity, and (2) support three-dimensional cell growth by providing a polymer matrix scaffold.^[1]^ These two requirements are fundamentally coupled, and in tension, in conventional nanoporous hydrogel networks. Hydrogel stiffness is tuned by increasing crosslink density and/or polymer concentration, which results in a mesh size of 5– 100 nm, orders of magnitude below the ∼10–20 μm scale of the cell body.^[2–4]^ In native tissues, cells circumvent this coupling by proteolytically remodeling the extracellular matrix (ECM), locally clearing space as they migrate and spread.^[5]^ In engineered hydrogels lacking relevant remodeling chemistry, however, the polymer network imposes persistent physical confinement, restricting spreading, protrusion extension, and migration regardless of bulk modulus.^[6,7]^ Macroporous hydrogels which contain larger pores on the scale of tens to hundreds of micrometers, address this limitation by providing pre-formed void space that decouples local confinement from bulk stiffness, enabling tissue-relevant moduli without trapping encapsulated cells^[8]^ Existing strategies to generate macroporosity, including porogen leaching,^[9]^ cryogelation,^[10]^ gas foaming,^[11]^ and granular hydrogel assembly,^[12]^ have advanced 3D cell studies both *in vitro*^[13]^ and *in vivo*,^[14]^ but share two limitations: they afford limited independent control over pore size and density, and most require fabrication conditions, e.g., freezing, extraction of sacrificial templates, or surfactant-stabilized foams, that preclude in situ cell encapsulation. Additionally, to our knowledge most macroporous platforms exhibit static bulk mechanical properties once formed, e.g. stiffness and crosslink density cannot be changed in situ, limiting the study of mechanically dynamic physiological processes that drive cell fate, migration, and tissue morphogenesis.^[15]^

How cells respond to these evolving cues, rather than static properties alone, has emerged as a central question in matrix biology. It is well established that the bulk elastic stiffness directly modulates cell phenotypes, from stem cell lineage specification^[16]^ to fibroblast and macrophage activation/ deactivation.^[17]^ Further work has established the important role of viscoelasticity and stress relaxation to modulate spreading, proliferation, and differentiation independently of elastic modulus.^[18–21]^ These responses unfold over hours to days, such that the trajectory of the mechanical environment, not simply its initial state, determines cell fate decisions. Probing how cells integrate and respond to stiffness cues requires dynamic biomaterials with programmable mechanical properties. ^[22,23]^

Photodynamic hydrogels represent a powerful solution to this material challenge, permitting user-controlled dynamicity in a hydrogel scaffold with spatiotemporal precision. Photodynamic materials incorporate photocleavable chromophores as crosslinkers within the polymer network, providing a direct handle to modulate network density and elastic stiffness.^[24]^ Irradiation at user-defined locations and doses has been used to generate stiffness gradients,^[25]^ channels,^[26]^ and patterned microenvironments,^[27]^ and to probe phenomena including mechanical memory, dynamic mechanotransduction, and their coupling.^[28]^ Yet despite the reported promise of photodynamic biomaterials, they are not broadly leveraged in the development of sophisticated tissue engineering platforms, likely due to the complex synthesis of common chromophores and the innate challenge of the nanoporous mesh inhibiting cellular responses in 3D. ^[29]^

With these design principles in mind, we sought to engineer a chemically simple polymer+crosslinker hydrogel platform capable of macropore formation and photosoftening. Our photocleavable crosslinker RuTetrazine leverages the well-established ruthenium(II) polypyridyl photochemistry for visible-light-triggered covalent bond cleavage,^[30,31]^ and incorporates two 1,2,4,5-tetrazine moieties for crosslinking with strained alkenes via inverse electron-demand Diels–Alder (iEDDA). iEDDA click chemistry is bioorthogonal and generates nitrogen gas as a byproduct, which becomes trapped in sufficiently dense polymer networks to form macroscopic pores (50-200 µm). To form a hydrogel, RuTetrazine was mixed at stoichiometric ratios with norbornene-modified hyaluronic acid (NorHA).^[32]^ We show that the elastic modulus of RuTetrazine-NorHA hydrogels can be tuned through incorporation of a photostable di-Tetrazine crosslinker and the size and density of macropores can be tuned through crosslink and polymer density. Importantly, we found that the novel RuTetrazine crosslinker is cytocompatible for long-term 3D cell culture, is non-mutagenic, and does not induce cell stress when incorporated into the hydrogel network. Finally, we wished to probe the effect of macropore presence and matrix softening on human mesenchymal stromal cells encapsulated within the HA polymer network and saw significant morphological changes in cells encapsulated in softened networks.

## 2. Results and Discussion

### 2.1. Synthesis and Characterization of RuTetrazine

RuTetrazine was prepared by reacting Ru(bpy)_2_Cl_2_·2H_2_O with AgOTf to generate the labile aquo intermediate, followed by coordination of 3-methyl-6-(3-pyridyl)-1,2,4,5-tetrazine and counterion exchange to the chloride salt (40% yield). The photostable diTetrazine and diTetrazine-modified MMP-RGDSK peptide crosslinkers were assembled through EDC/NHS activation of 3-(4-carboxyphenyl)-1,2,4,5-tetrazine, followed by conjugation with NH_2_-PEG_3_-NH_2_ or NH_2_-GPVGLIGGRGDSK-NH_2_ respectively. Norbornene-modified hyaluronic acid (NorHA) was synthesized by DMTMM-mediated amidation of sodium hyaluronate (90 kDa) with 5-norbornene-2-methylamine. All intermediates and final products were confirmed by ^1^H NMR spectroscopy, FTIR and mass spectrometry **(Figure S1-S9-S12)**.

Photoinduced ligand exchange of one PyrTet ligand was observed by UV-Vis absorbance spectroscopy **(Figure 1A)** and ^1^H NMR **(Figure S7)**. PyrTet ligand exchange with water was found to be rapid, requiring only 9J of energy (15 min irradiation with 10 mW blue light). Interestingly, despite the rapidity of ligand exchange, the quantum yield was observed to be quite low, only 0.048 ± 0.002.^[33]^ We hypothesize that this is due to the hydrophobic nature of the PyrTet ligand. To test this, we collected the quantum yield in a less polar solvent mixture (20%:80% methanol: water ratio), which increased the QY to 0.097 ± 0.025. We expect to observe rapid photocleavage when RuTetrazine is incorporated into a hydrogel network, which enhances the hydrophilicity of the ligand. ^[23,30]^ Prolonged irradiation (120 min, 450 nm, 10 mW) drove dissociation of the second 3-pyridine-tetrazine ligand, yielding the bis-aqua complex Ru(bpy)_2_(H_2_O) _2_ as evidenced by the emergence of a second isosbestic point at 485 nm (**Figure S10**). This stepwise photodissociation, also observed by ^1^H NMR (**Figures S7**), indicates differential photolability between the two coordination sites and allows selective access to either the mono- or bis-aquated product depending on irradiation dose.^[34]^ Photocleavage kinetics obeyed the Bunsen–Roscoe reciprocity law: cleavage curves collected at 10, 20, and 40 mW collapsed onto a single master curve when plotted against total light dose, reaching ∼100% cleavage at 2–3 J/cm^2^ regardless of incident power (**Figure S11**). This dose-dependent behavior allows irradiation time and intensity to be traded off interchangeably when designing photosoftening experiments.^[35]^

**Figure 1.**
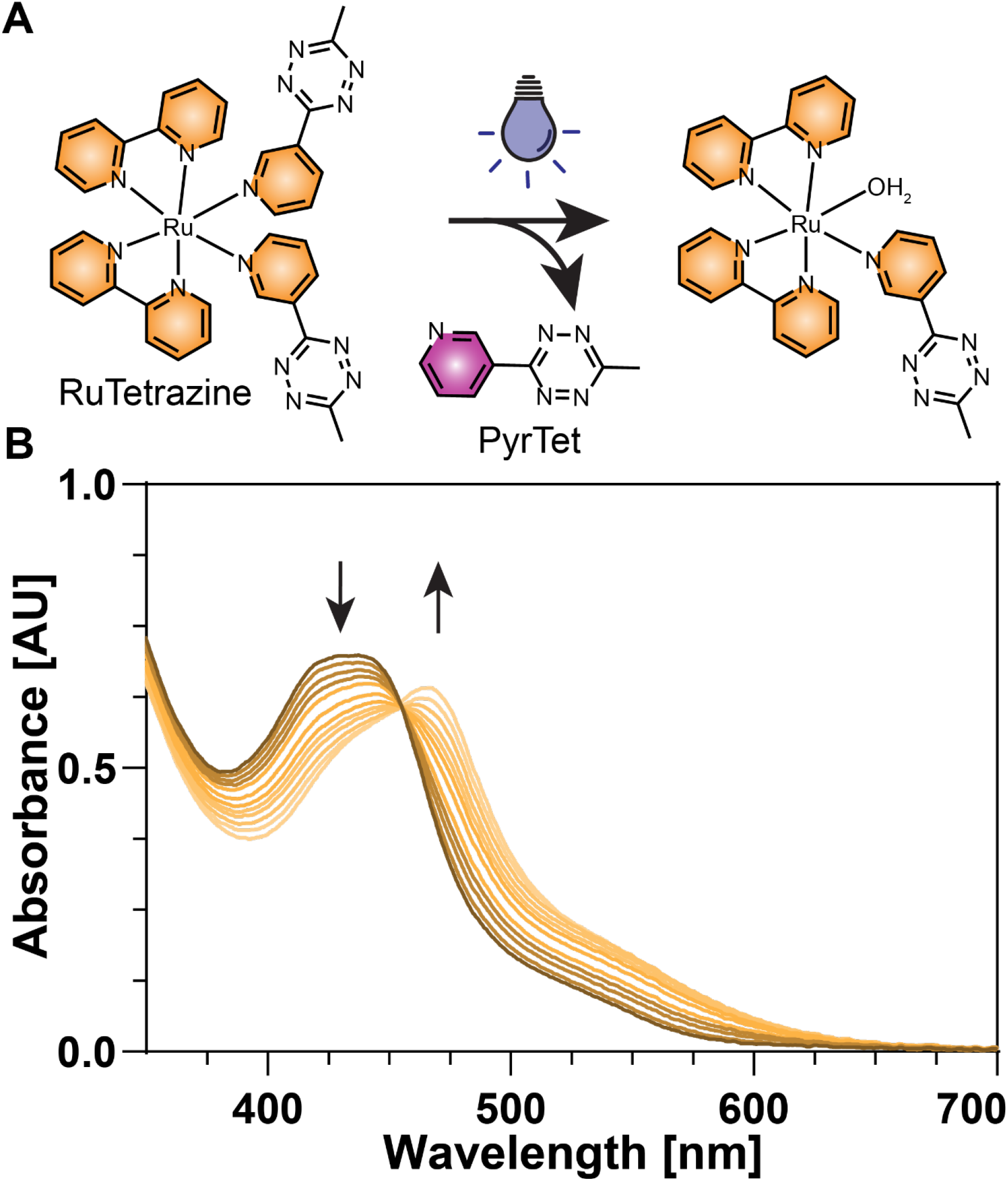
Photoinduced ligand exchange of RuTetrazine. A) RuTetrazine exchanges one pyridine ligand for a solvent molecule (water) under visible light irradiation (<550 nm). B) UV-Vis spectra tracking the photolysis in water (1.3 mM RuTetrazine, 450 nm, 10 mW) shows one isosbestic point at 455 nm indicating a clean A → B photolysis reaction with two absorbing species present.

### 2.2 Hydrogel formation

Inspired by the excellent photoresponse of RuTetrazine, we sought to generate a photosoftening hydrogel by mixing RuTetrazine with NorHA.^[36]^ Polymer network formation occurred through spontaneous crosslinking via iEDDA click reaction between tetrazine and norbornene (**Figure 2A**).^[32]^ This formulation allowed the formation of robust hydrogels from low-viscosity precursor solutions, ideal for future 3D cell encapsulation. To develop a framework of hydrogel formulation with this novel chemistry, we tested the elastic modulus of Ru-HA hydrogels with various polymer and crosslink concentrations. As expected, increasing the wt% of HA gave an increase in storage modulus from 1.5-3.5 kPa for 1 and 3 wt% HA respectively **(Figure 2B)**. We were able to achieve up to 90% modification of HA with norbornene groups via DMTMM activation, providing access to very stiff modulus approaching 10 kPa when mixed with precise 2:1 ratios of RuTetrazine crosslinker (e.g. 1 mol RuTetrazine for 2 mol norbornene groups) **(Figure 2C)**.^[37]^ With a large range of initial stiffnesses accessible in these Ru-HA hydrogels, we sought to develop a similar range of accessible intermediate stiffnesses following light irradiation and subsequent photocleavage of RuTetrazine crosslinks. To achieve controlled photosoftening rather than complete network dissolution, the photosensitive RuTetrazine crosslinker was combined with the photostable DiTet crosslinker within the same NorHA network **(Figure 2A)**. Photorheological measurements on pre-swollen hydrogels confirmed that the final post-irradiation stiffness is tunable through the photosensitive-to-photostable crosslinker ratio **(Figure 2D)**. For 3 wt% hydrogels, formulations with 80:20, 70:30, and 50:50 RuTetrazine:DiTet ratios softened by 87.3%, 76.3%, and 59.7%, respectively, with the 50:50 formulation retaining approximately 1.2 kPa storage modulus after irradiation **(Figure 2D, Table S1)**. Comparable tunable softening was observed for 2 wt% formulations, with initial stiffnesses approximately 33% lower than 3 wt% gels but similar percent reductions across crosslinker ratios (**Figure S14, Table S1)**. Hydrogel softening could be observed through visual inspection of the hydrogels: light-exposed softened hydrogels deepened in color, attributed to formation of Ru(bpy)_2_(PyrTet)(H_2_O) and appeared less structurally sound **(Figure 2E)**.

**Figure 2.**
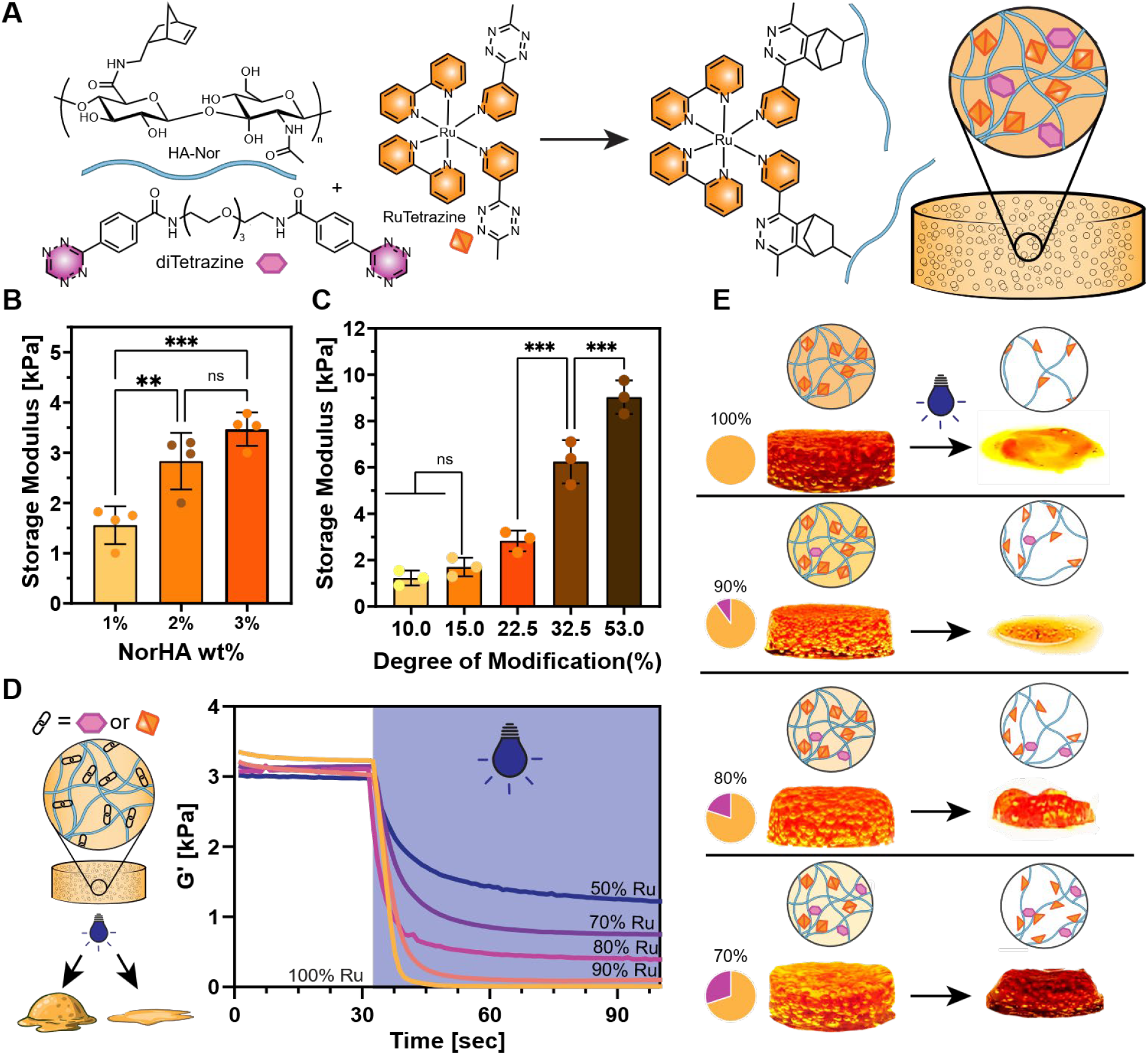
Mechanical Characterization of Ru-HA hydrogels. (A) Hydrogel formation via iEDDA crosslinking of NorHA with RuTetrazine and photostable DiTet; N_2_ evolved during gelation generates macropores in situ. (B) Storage modulus versus NorHA concentration. (C) Storage modulus versus degree of norbornene modification. (D)Photorheological characterization of 3 wt% hydrogels with varying RuTetrazine:diTet ratios under 455 nm irradiation (blue region, 20 mW). (E) Hydrogel appearance before and after irradiation at 70– 100% RuTetrazine content. Data: mean ± SD; **p < 0.01, ***p < 0.001.

Across all crosslinker ratios tested, hydrogels reached plateau storage modulus well within 20 minutes **(Figure S13)**, a critical window for maintaining consistent cell suspension during encapsulation. Although higher polymer concentrations (3 wt%) produced stiffer hydrogels, the rapid gelation kinetics posed practical challenges for cell encapsulation, including difficulty in casting uniform gels and potential mechanical stress on cells during mixing. We therefore selected a 2 wt% HA-Nor with 8.31 mM RuTetrazine, enough to fully react with 16.62 mM norbornene groups, approximating 35% modification of a 90 kDa HA, a formulation that balanced sufficient mechanical stiffness with manageable gelation times, reliable casting, and high cell viability.

Consistent with the iEDDA reaction mechanism, covalent network formation was accompanied by stoichiometric evolution of N_2_ gas. As described above, this intrinsic byproduct of iEDDA cycloaddition serves as the driving force for in situ macropore generation: in dense polymer networks, the evolved N_2_ becomes trapped during gelation, nucleating gas-filled cavities that constitute the macroporous architecture **(Figure 3A, S23, Supplementary Video 1)**. Macropore architecture was visualized in 1 wt%, 2 wt%, and 3 wt% RuTetrazine-crosslinked hydrogels stained with Cy5-tetrazine dye to visualize the polymer network. Pore diameters were quantified from the confocal images using ImageJ. **(Figure 3B, C)** Importantly, both the density and size distribution of these macropores are tuned by polymer concentration. Because crosslinker is added stoichiometrically to react with the available norbornene groups (1:1 Nor:Tet) at a fixed 35% degree of modification, increasing polymer concentration necessarily increases both the number of crosslinks and the total N_2_ evolved. These two contributions are mechanistically distinct: crosslinking generates the gas, while the polymer matrix captures and retains it as stable pores, but are tuned together. Increasing NorHA concentration from 1 to 3 wt% increased crosslink density and correspondingly the total volume of evolved N_2_, yielding a greater number of macropores with a significantly smaller average diameter: pore size decreased from 175 µm at 1 wt% to 85 µm at 2 wt% and 55 µm at 3 wt% (Figure 3C, p < 0.0001 across all pairwise comparisons). Lower-polymer formulations remain macroporous, producing fewer but larger pores, confirming that macroporosity does not require high polymer content. As macropore generation is tied to N_2_ bubble nucleation, we expected to observe a dense basement membrane-like polymer structure at the edge of the pores, which can be observed as halos in (**Figure 3B**, white arrows).

**Figure 3.**
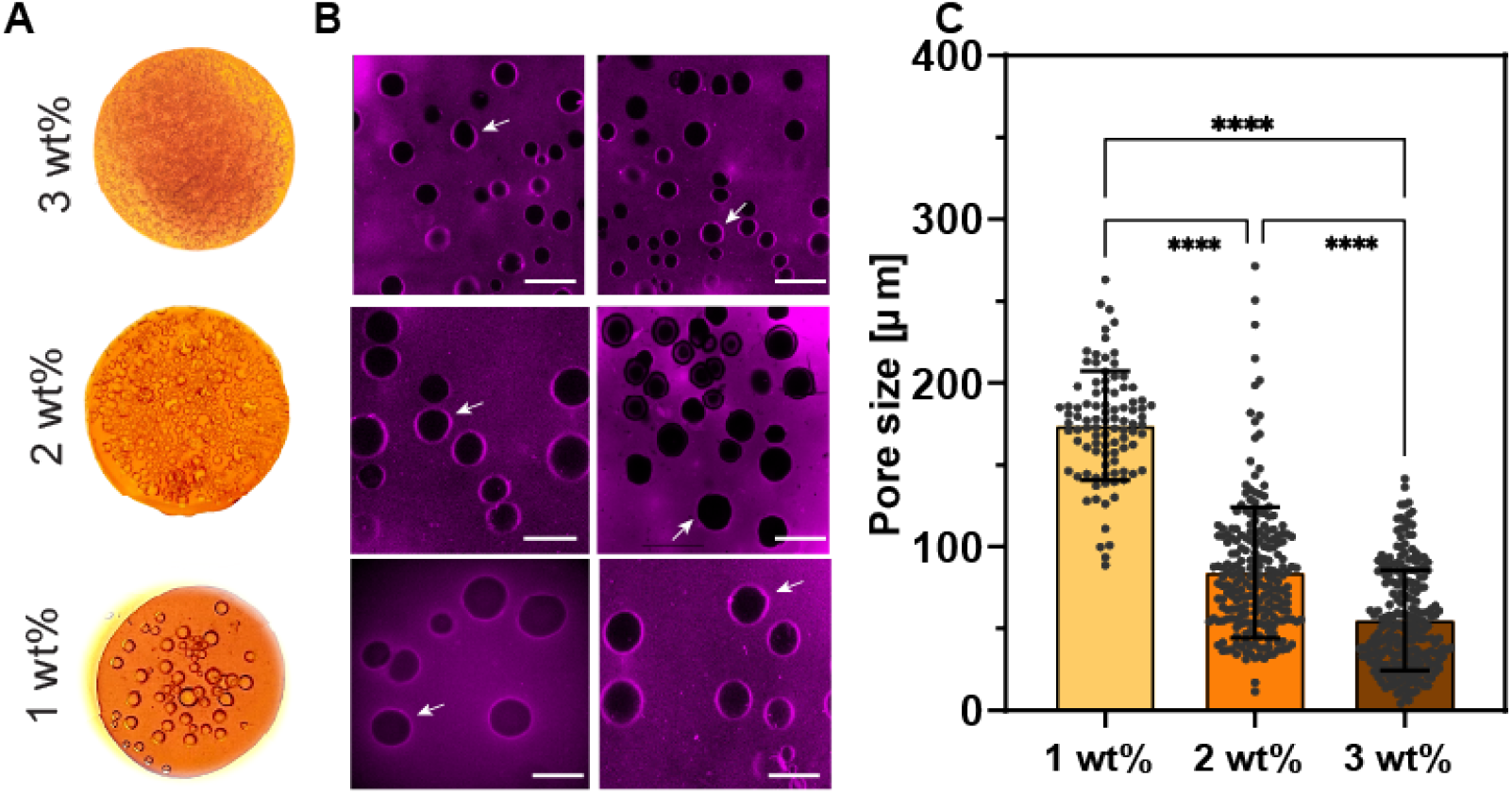
Macropore formation and diameter quantification. (A) Ru-HA hydrogels at 1–3 wt% with increasing bubble/pore density. Hydrogels were cast between glass slides. (B) Confocal fluorescence images of hydrogels at each concentration, revealing pore architecture. White arrows indicate dense basement membrane-like structures at the interface of macropores. Scale bars = 100 µm. (C) Quantification of pore size distributions across polymer concentrations. Data: mean ± SD; ****p < 0.0001.

### 2.3 Cytocompatibility of RuTetrazine and RuTet-NorHA Hydrogels

While ruthenium complexes possess excellent photochemical properties including radical-free covalent bond cleavage, cytocompatibility remains a significant concern facing their deployment as hydrogel crosslinkers in tissue engineering models. Previous work by our lab and others has confirmed the overall cytocompatibility of Ru-crosslinked hydrogels,^[30,38]^ in this work we sought to confirm that at our working range (4.15–8.31 mM) RuTetrazine is similarly non-toxic to encapsulated cells.

We first tested RuTetrazine’s IC50 value in normal human dermal fibroblasts (NHDFs) via WST-8 assay, a colorimetric assay for metabolic health, using standard protocols. Soluble RuTetrazine had an upper toxic limit of 0.27 ± 0.003 mM (**Figure S16**).^[39]^ While this is below the concentration used in Ru-HA hydrogels, our further investigation of cellular viability in 3D confirms significant toxicity modulation when RuTetrazine maintains bound to a polymer network (**Figure 4A,B**). Below the IC50 value, RuTetrazine was found to be non-mutagenic by Ames assay **(Figure S17)** and to not elicit a significant cellular ROS stress response **(Figure 4C,D)**.

**Figure 4.**
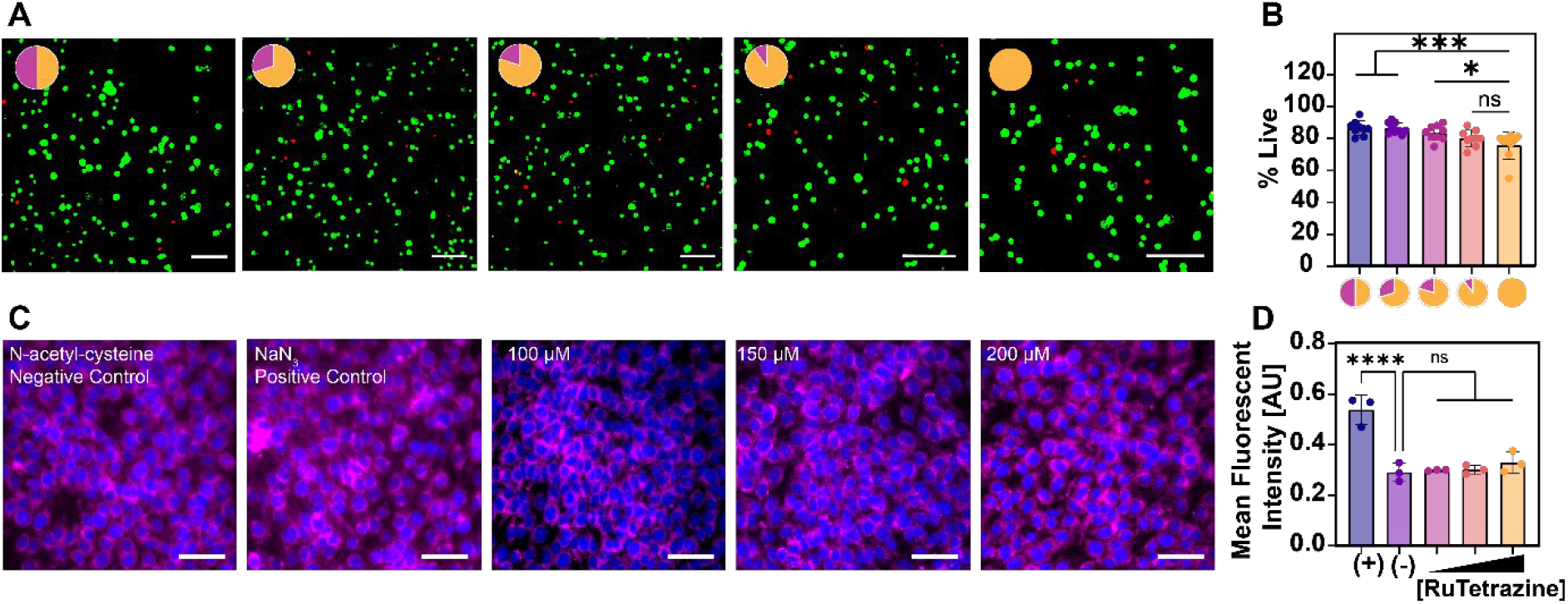
Cytocompatibility of RuTetrazine and Ru-HA Hydrogels. (A) Live/dead staining of NHDFs encapsulated in Ru-HA hydrogels (2 wt%, 35% norbornene modification) with varying RuTetrazine:DiTet ratios (50:50, 70:30, 80:20, 90:10, and 100:0), 24 h post-encapsulation. Calcein (green) indicates live cells; ethidium homodimer (red) indicates dead cells. Pie chart insets represent crosslinker composition (pink = RuTetrazine, orange = DiTet). (B) Live/dead image quantification shows >80% viability across all formulations. Scale bars = 100 µm. (C) CellROX Deep Red oxidative stress assay of NHDFs treated with negative control (N-acetyl-L-cysteine), positive control (sodium azide), and RuTetrazine at 100–200 µM. Hoechst (blue) marks nuclei; CellROX Deep Red (magenta) indicates intracellular ROS. (D) Fluorescence intensity quantification of shows RuTetrazine-treated cells exhibit ROS levels comparable to the negative control up to 200 µM. Scale bars = 100 µm. Data: mean ± SD; ****p < 0.0001, ***p < 0.001, *p < 0.05; ns = not significant.

In hydrogels, RuTetrazine was found to have a slight effect on final cell survival, assessed for NHDF and NIH-3T3 cells through live/dead cell staining using calcein green and ethidium homodimer. All formulations (50–100% RuTetrazine crosslinks) exhibited >80% viability 24 hrs post cell encapsulation **(Figure 4A, B, Figure S18)**. A slight, insignificant decrease in viability was observed with increasing RuTetrazine. This is attributed to the free RuTetrazine in the initial minutes of hydrogel formation prior to its sequestration in the polymer network. Post-irradiation (455 nm, 20 mW, 10 min) also exhibited >80% viability across all formulations, suggesting very little phototoxicity **(Figure S19)**. Encapsulated NHDFs showed ROS significantly lower than the sodium azide positive control, both before (**Figure S20**) and after photosoftening (**Figure S21**), confirming that neither encapsulation in Ru-HA hydrogels nor 455 nm irradiation induces measurable oxidative stress or affects overall viability.This result confirms our hypothesis that if the RuTetrazine is sequestered in the polymer network, this will modulate its toxicity and significantly increase its biocompatibility, due to the main mechanism leading to cell death, which is DNA binding and stress-related apoptosis. ^[40]^

### 2.4 Cellular Response to Photosoftening Macroporous Hydrogels

Having established the unique polymer network structure surrounding macropores in this material formulation, we sought to further probe the cellular microenvironment in the bulk and pore-adjacent regions of the hydrogel. A critical feature of this system is that the N_2_-filled bubbles fill upon equilibration and swelling in medium, converting gas-phase voids into media-saturated macropores accessible to cells. Images of 1–3 wt% hydrogels incubated in DMEM over 10 days captured this transition, with visible bubbles diminishing between Day 1 and Day 10 of culture (**Figure S24**).

To explore the cellular microenvironment around the in situ bubbles, NHDFs were encapsulated into the bulk hydrogel during casting. Confocal imaging of encapsulated NHDFs 24 hrs after encapsulation revealed that cells were dispersed throughout the bulk hydrogel, with those cells near bubble edges entrapped in the tight, aligned polymer network near bubble edge (**Figure 5**). This edge-localized distribution reflects a passive, fabrication-driven process: as N_2_ bubbles expand during gelation, they compress the surrounding polymer into a dense, basement-membrane-like shell at the pore wall **(Figure 3B-Figure S22)** and mechanically squeeze nearby cells into alignment around the void. This basement-membrane-like interface is a distinctive feature of this macroporous system relative to other 3D scaffolds. ^[41]^

**Figure 5.**
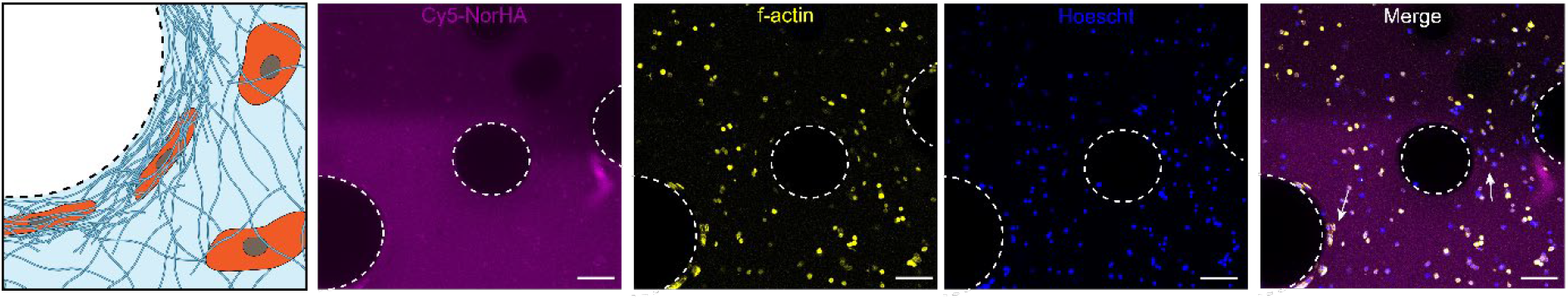
Bubble entrapment aligns and concentrates HA polymers around hydrogel-pore interface. NHDFs encapsulated in 2 wt% hydrogels (35% norbornene modification) and imaged for NorHA network (magenta), cytoskeleton (yellow), and nucleus (blue). Cells are elongated in the immediate vicinity of pores due to polymer network compression from bubble formation and entrapment (white arrows). Scale bars = 200 µm.

Intrigued by the different microenvironments present in our hydrogel platform, we sought to probe the impact of matrix softening on cell morphology and spreading. To this end, we encapsulated hMSCs in Ru-HA hydrogels (2 wt% NorHA, 25% norbornene modification). For these experiments, the photostable diTet crosslinker was replaced with an MMP-cleavable ditetrazine peptide bearing pendant RGD sequences, enabling both integrin-mediated adhesion and cell-directed matrix remodeling within the network (**Figure 6A**).^[42]^ Photorheological characterization confirmed that substituting the MMP-degradable RGD-bearing ditetrazine peptide for DiTet preserves the photosoftening response: 2 wt% NorHA hydrogels formulated with an 80:20 RuTetrazine:Ttz-MMP-RGDSK-Ttz crosslinker ratio softened from an initial storage modulus of 2.27 kPa to 0.54 kPa (∼76% reduction) following 5 min of 455 nm irradiation at 20 mW (**Figure S15**), comparable to the softening behavior observed in diTet-crosslinked networks (**Figure 2D, Figure S14**).

**Figure 6.**
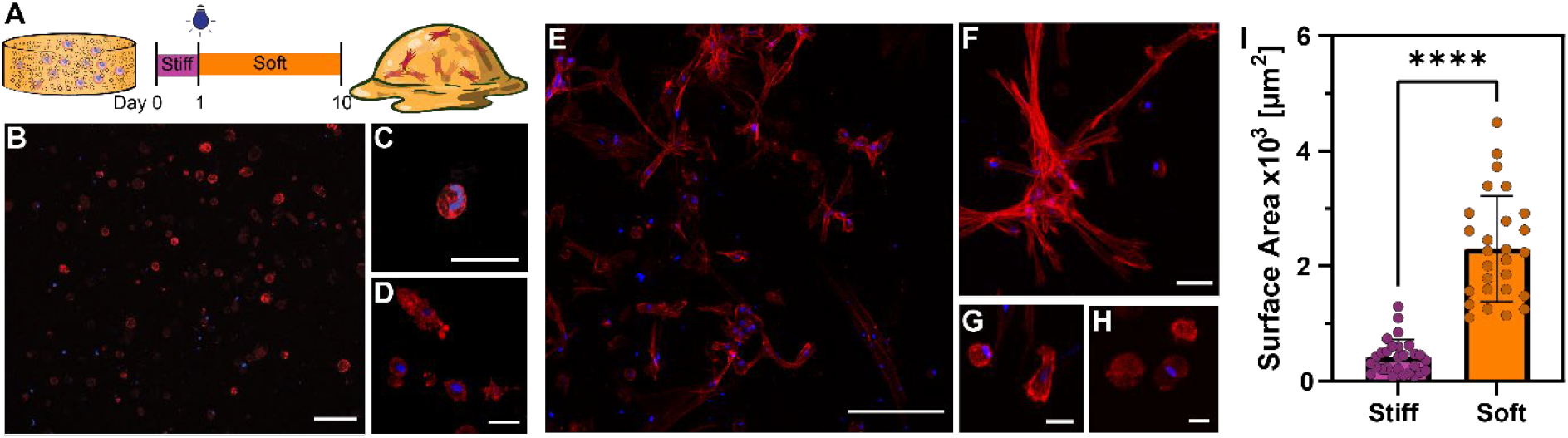
Photosoftening drives hMSC spreading in macroporous Ru-HA hydrogels. (A) hMSCs were encapsulated at Day 0, allowed to equilibrate for 24 hours, then photosoftened on Day 1 (455 nm, 20 mW, 10 min) or maintained in non-irradiated control conditions; cultures were imaged at Day 10. (B–D) Representative fluorescence micrographs of hMSCs in non-photosoftened (stiff) hydrogels at Day 10, showing predominantly rounded morphology (B: field view, Scale bar = 200 μm; C, D: single-cell insets, scale bar 50 μm). (E–H) Representative fluorescence micrographs of hMSCs in photosoftened hydrogels at Day 10, showing extensive cell spreading, protrusion extension, and formation of interconnected multicellular networks (E, F: field views, scale bar = 200 μm; G, H: single-cell insets, scale bar = 50 μm). F-actin (red, phalloidin) and nuclei (blue, Hoechst). (I) Quantification of cell surface area in photosoftened (orange) versus non-photosoftened (purple) conditions. Photosoftened cells exhibited an approximately 6-fold increase in surface area relative to non-photosoftened controls. Data: mean ± SD; ****p < 0.0001.

To probe the cellular response to photosoftening, hydrogels were cast with hMSCs, allowed to gel and equilibrate for 24 hours, then photosoftened *in situ* by irradiation with 455 nm light (20 mW, 10 min) and maintained in culture for an additional 9 days (**Figure 6B**). In non-irradiated, stiff hydrogels, cells remained predominantly rounded throughout the 10-day culture period, with minimal protrusion formation and a compact spherical morphology characteristic of cells confined within a stiff matrix (**Figure 6C**). In striking contrast, cells encapsulated in photosoftened hydrogels underwent extensive spreading, extending elongated protrusions and forming interconnected multicellular networks that spanned the macroporous architecture by Day 10 (**Figure 6D**). Quantification of cell surface area revealed an approximately 6-fold increase in photosoftened conditions relative to non-photosoftened controls (∼6,500 μm^2^ vs ∼1,100 μm^2^, p < 0.0001; **Figure 6E**).^[5]^ This dramatic morphological divergence from a common starting population demonstrates that the matrix softening drives the observed spreading response. This finding is consistent with prior demonstrations that 3D matrix mechanosensing in HA hydrogels depends on accessible adhesive and mechanical cues rather than encapsulation conditions alone.^[43]^ Together, these results show that photosoftening, by cleaving RuTetrazine crosslinks to reduce matrix stiffness and enlarge mesh size, enables encapsulated cells to spread extensively in 3D. The incorporation of an MMP-degradable, RGD-bearing crosslinker further allows cells to engage and remodel the surrounding network, supporting the elongated, interconnected morphologies observed after softening.

## Conclusion

This report describes the design of the first macroporous, photosoftening hydrogel for tissue engineering. This hyaluronic acid-based hydrogel combines in situ macropore generation with visible-light photosoftening through a single crosslinking chemistry. The photocleavable RuTetrazine crosslinker forms hydrogels with norbornene-modified hyaluronic acid via iEDDA click chemistry, and both initial stiffness and the extent of softening are tunable through polymer concentration and the photosensitive-to-photostable crosslinker ratio. The crosslinker is non-mutagenic and does not elicit oxidative stress once network-bound, supporting long-term cell survival with minimal sub-toxic effects.

The unique combination of a photocleavable crosslinker and macroporous structure offers a way to independently tune porosity, which is inherently coupled with polymer concentration, and final material stiffness. While a high concentration of both polymer and crosslinker are necessary to achieve the most porous structure, subsequent photosoftening after complete gelation provides an opportunity to decouple material stiffness and macroporosity, which we showed by encapsulated hMSCs in the Ru-HA hydrogel formulated with an MMP-RGD peptide crosslinker. Encapsulated hMSCs exhibited significant morphological response to photosoftening, with an approximately 6-fold increase in surface area relative to non-softened controls.

The single-step encapsulation, macropore generation, and spatiotemporal control over stiffness and crosslink density will enable further studies in cell-interface interaction, a more accurate platform to model basement membrane structure, and spatiotemporal effects of matrix softening on cell morphology, differentiation, and growth.

## Acknowledgements

The authors thank Adam Fries (Genomics and Cell Characterization Core Facility, University of Oregon) for assistance with confocal and widefield fluorescence microscopy and image analysis.

## Data Availability Statement

The authors encourage interested readers to contact Dr. Rapp for access to supplementary information and synthetic procedures.

