## Supplementary Information for "Photosoftening Macroporous Hydrogels for Dynamic Tissue Engineering"

**Materials**

Sodium hyaluronate was purchased from Lifecore Biomedical (HA60K-5, 66K-99) 5-Norbornene-2-methylamine (CAS 95-10-3), EDC·HCl (CAS 25952-53-8), hydrazine monohydrate (CAS 7803-57-8), potassium hexafluorophosphate (CAS 17084-13-8), *N*-hydroxysuccinimide (≥98.0%, CAS 6066-82-6), *N*,*N*-diisopropylethylamine (CAS 7087-68-5), and DMTMM (CAS 3945-69-5) were purchased from TCI. 3-Pyridinecarbonitrile (98%), silver trifluoromethanesulfonate (CAS 2923-28-6), elemental sulfur (CAS 7704-34-9), and the WST-8 reagent (≥97% HPLC, CAS 193149-74-5) were purchased from Sigma-Aldrich. *cis*-Bis(2,2′-bipyridine)dichlororuthenium(II) dihydrate (CAS 15746-57-3) was purchased from Beantown Chemical, 4-cyanobenzoic acid (CAS 619-65-8) from Alfa Aesar, amino-PEG$_3$-amine from BroadPharm, and Cy5-tetrazine from Lumiprobe. The 5041-2BS Ames MOD-ISO kit was used for Ames assay. NH2-GPVGLIGGRGDSK-NH$_2$ was custom-synthesized by GenScript. Normal human dermal fibroblasts (NHDFs) and NIH/3T3 mouse embryonic fibroblasts were purchased from ATCC. Human mesenchymal stromal cells (hMSCs) from RoosterBio. For cell culture and staining, phenol red–free DMEM/F-12 (cat. 21041025), CellROX Deep Red reagent (C10422), calcein AM, ethidium homodimer-1 (E1169), Hoechst 33342 (H3570), Alexa Fluor 488–phalloidin, and Alexa Fluor Plus 647–phalloidin were purchased from Invitrogen/Thermo Fisher Scientific. Fetal bovine serum (FB72) was purchased from CellPro and Rooster Nourish medium (K82003) from RoosterBio. All solvents were used as received unless otherwise noted.

**Instrumentation**

Rheological and photorheological measurements were performed on a Discovery Hybrid Rheometer HR-30 (TA Instruments) operated with TRIOS software, using a quartz insert for in situ photorheology. Confocal fluorescence imaging was performed on a Zeiss LSM 880 confocal microscope and widefield fluorescence imaging on a Leica TCS SPE microscope. $^1$H NMR spectra were acquired on Bruker spectrometers (500 and 600 MHz). High-resolution ESI mass spectra were collected at the University of Chicago (Noyes Laboratory mass spectrometry facility). FTIR spectra were acquired on a Shimadzu IRAffinity-1S spectrometer equipped with a QATR single-reflection diamond ATR accessory. UV-Vis absorbance spectra acquired on Thermo Scientific NanoDrop One. Visible-light irradiation for photolysis and photosoftening was delivered by a collimated 450/455 nm Mic-LED (Prizmatix, Southfield, MI, USA) driven by a BLCC-04 current controller**.** Image analysis was performed in Fiji/ImageJ and Imaris; IC50 fitting used the AAT Bioquest calculator (four-parameter logistic model); statistics were performed in GraphPad Prism.

**Methods**

**Synthesis of 3-(4-carboxyphenyl)-1,2,4,5-tetrazine**

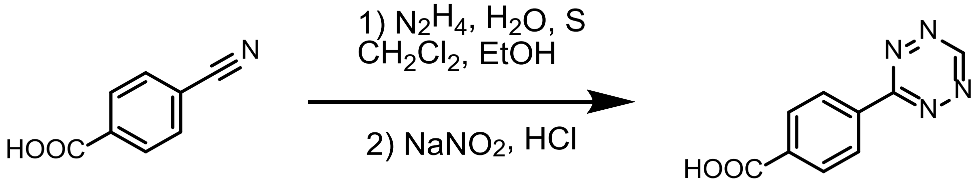

In, 4-cyanobenzoic acid (147.13 mg, 1.0 mmol, 1.0 eq, MW: 147.13 g/mol), dichloromethane (63.70 µL, 1.0 mmol, 1.0 eq, MW: 84.93 g/mol), ethanol (1 ml), and elemental sulfur (64 mg, 2.0 mmol, 2.0 eq, MW: 32.06 g/mol) was combined in a 30 mL microwave reaction vessel. While stirring, hydrazine monohydrate (0.4 mL, 8.0

mmol, 8.0 eq, MW: 50.06 g/mol) was added dropwise. The vessel was sealed and heated to 50°C overnight. (WARNING: Exercise caution as sealed vessels develop significant pressure when heated, especially during scale-up procedures). After 18 hours, dichloromethane (3 mL) was added to the reaction mixture, followed by a solution of sodium nitrite (0.69 g, 10.0 mmol, 10.0 eq, MW: 68.995 g/mol) dissolved in water (10 mL). Glacial acetic acid (3.42 mL, 60.0 mmol, 60.0 eq, MW: 60.05 g/mol) was added dropwise, resulting in the formation of an intense red coloration. The reaction mixture was extracted with large excess of ethyl acetate. The organic phase was dried over anhydrous magnesium sulfate ($MgSO_4$), filtered and concentrated under reduced pressure. [1] The product was purified by recrystallization from ethyl acetate with 3% acetic acid as a red solid (70% yield). $^1H$ NMR (500 MHz, DMSO-$d_6$) δ 10.66 (s, 1H), 8.62 (d, 2H), 8.22 (d, 2H) **(Figure S3).**

**Synthesis of Tetrazine-NHS**

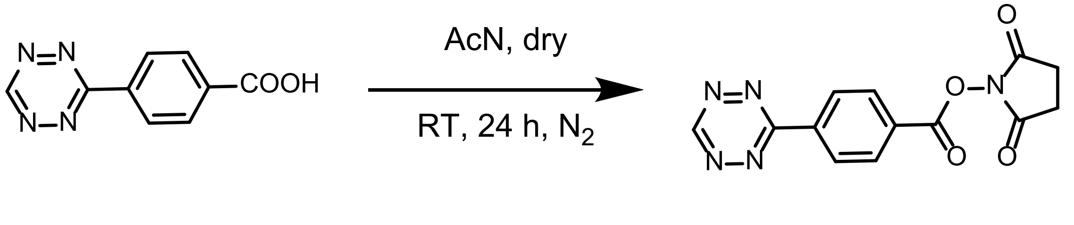

3-(4-Carboxyphenyl)-1,2,4,5-tetrazine (0.068 g, 0.34 mmol, 1.0 equiv), *N*-hydroxysuccinimide (0.060 g, 0.52 mmol, 1.1 equiv), and *N*-(3-dimethylaminopropyl)-*N*′-ethylcarbodiimide hydrochloride (EDC·HCl, 0.097 g, 0.51 mmol, 1.1 equiv) were dissolved in anhydrous acetonitrile (2 mL) under $N_2$. The reaction was stirred overnight at room temperature, after which the solvent was removed under reduced pressure. The residue was resuspended in ethyl acetate (10 mL), and the organic layer was washed with water (3 × 10 mL), dried over $MgSO_4$, filtered, and concentrated under reduced pressure to afford the crude product as a pink solid. The crude material was dissolved in a minimum volume of anhydrous dichloromethane (~1 mL per 100 mg; anhydrous acetonitrile may be substituted) and added dropwise, with swirling, to cold diethyl ether (−20 °C, 10–20 volume equivalents) in a centrifuge tube. The resulting pink precipitate was isolated by centrifugation, the supernatant decanted, and the pellet washed with cold diethyl ether (2 × 10 mL). The pellet was dried under vacuum at room temperature (2–4 h) to give TTZ-NHS ester as a pink solid (64 % yield).[2] $^1H$ NMR (500 MHz, DMSO-$d_6$): δ 8.80 (s, 2H, tetrazine-H), 8.45 (d, 2H, Ar-H), 8.25 (d, 2H, Ar-H), 2.92 (s, 4H, NHS-$CH_2$). **(Figure S4).**

**Synthesis of Tetrazine-PEG3-Tetrazine (diTet)**

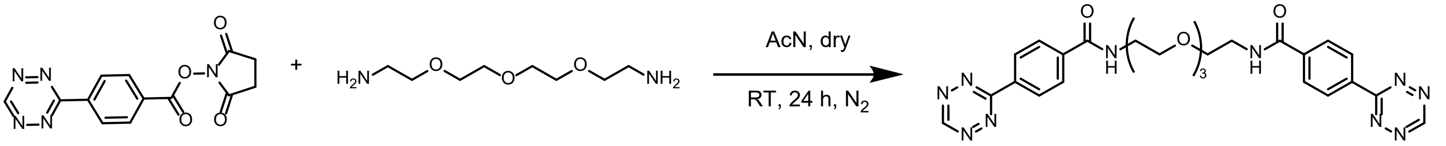

TTZ-NHS ester (20 mg, 0.067 mmol, 2.1 eq, MW: 299.25 g/mol) was dissolved in dry DMF (3.5 mL) under an inert atmosphere. DIPEA (16.7 µL, 0.096 mmol, 3.0 eq, MW: 129.24 g/mol, d = 0.742 g/mL) followed by Amino-PEG3-Amine (6.12 mg, 0.032 mmol, 1.0 eq, MW: 192.26 g/mol; alternatively, 5.96 µL, d = 1.027 g/cm$^3$) were added to the stirring solution and the reaction mixture stirred at room temperature for 18 hours. The use of anhydrous conditions is critical as NHS esters are moisture-sensitive and readily hydrolyze in the presence of water. The product was purified by precipitation from cold diethyl ether (-20°C, 65 mL, 10-fold excess volume) followed by dialysis against ultrapure water for 48 hours with regular water changes to remove small molecule impurities. Product was lyophilized and stored as a powder.[3] $^1H$ NMR (500 MHz, DMSO-$d_6$) δ 10.62 (s, 2H, tetrazine-H), 8.65 (d, 4H, Ar-H), 8.10 (d, 4H, Ar-H) **(Figure S5)**.

#### Synthesis of Ttz-MMP-RGDSK-Ttz Peptide Crosslinker

A foil-wrapped, oven-dried 4 mL vial equipped with a magnetic stir bar was charged with the MMP-cleavable RGD-bearing peptide ($NH_2$-GPVGLIGGRGDSK-$NH_2$; MW: 1212.36 g/mol, 20 mg, 16.5 µmol, 1.0 eq) under a nitrogen atmosphere. Anhydrous DMF (800 µL) was added, and the mixture was stirred for 2–5 min at room temperature until fully dissolved. (Note: if peptide solubility in DMF is poor, anhydrous DMSO may be substituted.) DIPEA (MW: 129.24 g/mol, d = 0.742 g/mL, 28.7 µL, 21.3 mg, 165 µmol, 10 eq) was added, and the solution was stirred for an additional 2 min. In a separate dry vial, TTZ-NHS (MW: 299.25 g/mol, 29.6 mg, 99 µmol, 6.0 eq) was dissolved in anhydrous DMF (300 µL) immediately before use, yielding a bright pink-red solution. The TTZ-NHS solution was added dropwise to the stirring peptide mixture, and the combined reaction was wrapped in foil and stirred under $N_2$ at room temperature for 16 h. The crude reaction mixture was diluted with water containing 0.1% formic acid, filtered, and purified by preparative reverse-phase HPLC on a C18 column (250 × 21.2 mm) at a flow rate of 15 mL/min (A = 0.1% FA in $H_2O$; B = 0.1% FA in MeCN), with detection at 220 nm and 525 nm. The gradient was held at 5% B for 5 min, then ramped linearly to 40% B at 15 min and to 95% B at 20 min, and held at 95% B until 30 min. The bis-modified product eluted latest (≈20 min), after the singly-modified intermediate (≈15 min) and well after the unmodified peptide (≈10 min) and was identified by its characteristic deep pink color. Pink fractions were pooled, diluted with water to <40% MeCN, frozen at –80 °C, and lyophilized to afford Ttz-MMP-RGDSK-Ttz as a deep pink powder. The purified product was stored at –80 °C in a foil-wrapped, argon-flushed cryovial and used within 1–2 months to minimize degradation of the H-tetrazine moieties.[3] HRMS (ESI) m/z: $[M+2H]^{2+}$ calcd for $C_{69}H_{99}N_{25}O_{19}^{2+}$, 790.8774; found, 790.8788. **(Figure S9)**

#### Synthesis of NorHA

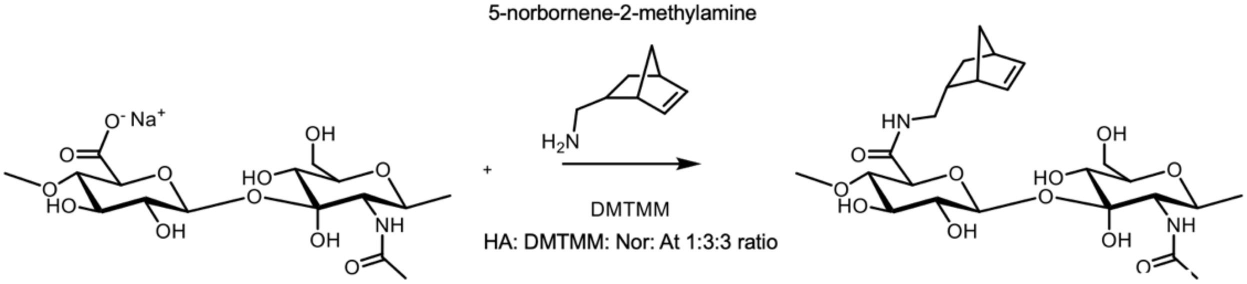

Sodium hyaluronate (MW: 90 kDa, 0.4 g, 5.28 × $10^{-4}$ mol of repeating units, 1 eq) was fully solubilized in 40 mL of MES buffer at pH 5.5 to achieve a 1% w/v solution. DMTMM (MW: 276.72 g/mol 0.83 g, 1.50 × $10^{-3}$ mol, 3 mol eq) was introduced and the mixture stirred for 10 minutes. Subsequently, 5-norbornene-2-methylamine (MW: 123.20 g/mol, 0.256 mL, ~1.06 × $10^{-3}$ mol, 2 mol eq) was added dropwise, and the reaction proceeded at room temperature for 24 hours to achieve a target degree of modification between 20-40%, which was found optimal for subsequent applications. For product isolation, saturated NaCl solution (16 mL, corresponding to 0.4× the MES volume) was added with 20 minutes of stirring, followed by gradual precipitation over approximately 3 hours using 200 proof ethanol (100 mL, 2.5× the MES volume). The precipitated product was collected through vacuum filtration and rinsed with both 140 and 200 proof ethanol. The material was then redissolved in deionized water and subjected to dialysis using 6-8 kDa molecular weight cutoff tubing in Milli-Q water for 72 hours, with water replacement every 3 hours using a 3L beaker maintained at room temperature.[4] The purified product was subsequently lyophilized and stored at -20°C in powder form.[5] H NMR (500 MHz, $D_2O$) δ 6.20–5.90 (m, 2H, norbornene vinyl -CH=CH-), 4.50–3.20 (m, 10H, HA sugar ring), 2.90–2.70 (m, 2H, $-CH_2-NH-$), 2.30–0.50 (m, norbornene ring protons). **(Figure S6, S12)**

##### Quantum Yield Determination

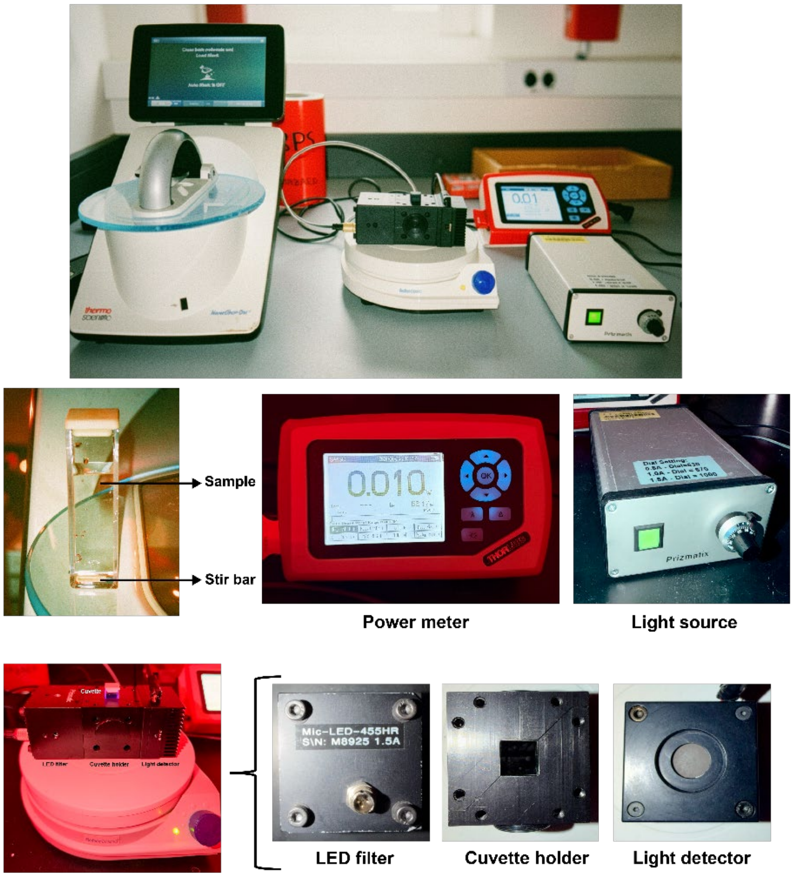

Quantum yield was determined by the following equation:

$$\frac{\frac{d[RuTetrazine]}{dt}}{\frac{d(photons)}{dt}} = Quantum\ Yield$$

Where

$$\frac{d[RuTetrazine]}{dt} = k_{obs}[RuTetrazine]_i$$

$k_{obs}$ is the observed rate constant for the pseudo-first order photochemical reaction.
And

$$\frac{d(photons)}{dt} = \frac{Power}{E_{ph}}$$

Where $E_{ph}$ is the energy of the photon, defined as

$$E_{ph} = \frac{hc}{\lambda}$$

And *Power* is the power of incident light.

The observed rate constant $k_{obs}$ was determined by kinetic fitting of photolysis UV-Vis data (Abs@photoproduct vs. time) to an equation of the form:

$$y = Ae^{-\frac{x}{\tau}} + y_0$$

Where $y$ is the absorbance at the photoproduct $\lambda_{max}$ (470 nm) and $x$ is time. $y_0$ is the molar absorptivity of the photoproduct times the concentration of the photoproduct, approximated as the final absorbance value following complete photolysis. $\tau = 1/k_{obs}$.

**Bubble Size Quantification in Macroporous Ru-HA Hydrogels**

Ru-HA hydrogels (50 µL total volume) were prepared at 1, 2, and 3 wt% with 35% norbornene modification according to the procedure above. To visualize the polymer network, NorHA was pre-reacted with Cy5-tetrazine dye (Lumiprobe) prior to hydrogel casting: a 5 mM solution of Cy5-tetrazine in PBS was combined with NorHA and allowed to react via iEDDA click chemistry with a small fraction of the pendant norbornene groups, producing fluorescently labeled NorHA stock. The labeled NorHA was then used directly in hydrogel formulation as described above, ensuring uniform incorporation of the dye throughout the polymer network during gelation. Fluorescently labeled hydrogels were imaged on a Leica widefield microscope using a 10X objective with a Cy5 filter set. Complementary images were acquired on a Zeiss LSM 880 confocal microscope using identical acquisition settings to confirm pore architecture at higher resolution. Z-stack images were acquired at a single representative region per hydrogel to confirm that the full diameter of each bubble was captured in the focal plane, ensuring accurate diameter measurements. Bubble diameters were quantified from fluorescence images using Fiji software (**Figure S23**).

**Rheometric Analysis**

Rheological measurements were conducted at 25 °C on a TA Instruments rheometer operated with TRIOS software using an 8 mm parallel plate geometry. Oscillatory time sweep experiments were performed at a frequency of 1 Hz and a strain amplitude of 1%, within the linear viscoelastic region of the hydrogels, to determine the storage (G′) and loss (G″) moduli. These parameters were selected to ensure non-destructive testing while providing reliable mechanical characterization of the crosslinked networks. For photorheology, a quartz stage coupled to a visible light source by fiber optic cable enabled measurement of rheological properties during gelation and light irradiation of the hydrogels.

**Photorheological Characterization of Ru-HA Hydrogels with Tunable Photosoftening**

50 µL hydrogels were allowed to gel at 37 °C for 30 minutes, and swelled in PBS overnight at 37 °C. Swollen hydrogels were then loaded onto a TA after establishing a baseline storage modulus (G′), hydrogels were irradiated in situ with 455 nm light (20 mW) for 5 minutes to induce photocleavage of RuTetrazine crosslinks, while the storage modulus was continuously monitored before, during, and after irradiation to capture the full photosoftening response. Overall, the extent of photosoftening was inversely correlated with photostable crosslinker content, with higher DiTet ratios requiring extended irradiation times **(Figure S13-15-Table S1).**

**Table S1. Pre- and Post-Photosoftening Storage Moduli**

| 3% gel Crosslinker ratio | Initial stiffness | Stiffness after photosoftened |
|---|---|---|
| **80** | 3079 | 391 |

| | | |
|---|---|---|
| **70** | 3150 | 748 |
| **50** | 3016 | 1216 |
| **2% gel Crosslinker ratio** | Initial stiffness | Stiffness after photosoftened |
| **80** | 2048 | 328 |
| **70** | 2117 | 849 |
| **50** | 2111 | 1031 |

**Cell Culture and Maintenance**

Normal human dermal fibroblasts (NHDFs) were cultured in T75 flasks using phenol free Dulbecco's Modified Eagle Medium (Thermo Scientific #21041025) supplemented with 10% fetal bovine serum and 1% penicillin-streptomycin. Human mesenchymal stromal cells (hMSCs) were cultured in T75 flasks prior to encapsulation using Rooster Nourish (Rooster Bio, K82003) complete media kit. Cells were maintained at 37°C in a humidified atmosphere containing 5% carbon dioxide. Upon reaching 80-90% confluency, cells were washed with PBS, lifted using TrypLE, and counted for accurate cell seeding. All experiments were performed using cells between passages 5 and 12.

**$IC_{50}$ Determination**

$IC_{50}$ values for RuTet and diTet crosslinkers were determined using a WST-8 metabolic assay. Normal human dermal fibroblasts (NHDFs, passages 5–12) were cultured according to the protocol above. Stock solutions of both the photosensitive ruthenium-di-tetrazine crosslinker and the photostable control crosslinker were prepared at 1 mM concentration in sterile PBS (pH 7.4), sterile-filtered through 0.22 μm membranes, and stored protected from light at -25°C. Working dilutions were prepared fresh in complete culture medium immediately prior to treatment. For cytocompatibility assessment, NHDFs were seeded into 96-well tissue culture plates at a density of 7,500 cells per well and allowed to adhere overnight. Culture media was then replaced with a media containing crosslinker at concentrations of 0, 50, 150, 250, 350, 450, 550, 650, and 750 μM, and cells were incubated for 24 hours under standard culture conditions. Both the photosensitive and photostable crosslinkers were evaluated on separate plates, with untreated cells serving as controls. Following the treatment period, treatment media was aspirated and wells were washed with PBS to remove excess crosslinker. Then, 100 μL of WST-8 and mPMS reagent were added to each well, and the plates were incubated for 3 hours. Absorbance was measured at 450 nm using a microplate reader, and viability was expressed as a percentage relative to untreated controls. Half-maximal inhibitory concentration ($IC_{50}$) values were determined from dose-response curves using the AAT Bioquest IC50 calculator **(Figure S16).**

**AMES assay for RuXlinker**

Mutagenicity potential of the photodynamic crosslinker was determined via the 5041-2BS Ames MOD-ISO Bacterial Strain Kit (EBPI Toxicity Tests) through exposure of increasing log concentrations of Ru-Tet to auxotrophic histidine *Salmonella typhimurium* mutants in histidine deficient culturing conditions. Carcinogenic compounds cause frameshift and or point mutations which revert the strain back to a prototrophic state of histidine production allowing for detectable colony formation. The salmonella strain is pre exposed for 100 minutes in triplicate before being transferred to a reversion solution and plated onto a 96 well plate prior to a 3 days incubation. The number of colonies are counted via a colorimetric analysis where yellow indicates significant histidine reversion (colony growth) and light purple indicating no or low histidine reversion (colony inhibition) **(Figure S17).**

**Oxidative Stress Assessment of Soluble RuTetrazine by CellROX Deep Red Assay**

Normal human dermal fibroblasts (NHDFs, passages 5–12) were seeded at 15,000 cells/cm² in 12-well plates (52,500 cells/well) in 1 mL complete media and allowed to adhere overnight. On Day 2, media was aspirated and replaced with fresh media containing RuTetrazine at final concentrations of 25, 50, 100, 150, or 200 µM and incubated overnight. A negative control was prepared by treating cells with 5 mM N-acetyl-L-cysteine (NAC; MW 163.19 g/mol, dissolved in warm media) for 24 hours to suppress basal ROS levels. A positive control was prepared by aspirating media and treating cells with 200 µM sodium azide ($NaN_3$; MW 65.01 g/mol, dissolved in PBS) for 2 hours to induce oxidative stress without cytotoxicity. On Day 3, treatment media was aspirated and wells were washed once with PBS. A staining solution was prepared in warm media containing CellROX Deep Red (5 µM), Hoechst 33342 (5–10 µg/mL), and Calcein AM (3.5 µM). Staining solution was incubated with all cells for 30 minutes at 37 °C. Following incubation, the dye suspension was aspirated and cells were washed three times with PBS. Live cells were imaged immediately on a Leica widefield microscope using a 20X objective, with DAPI and Cy5 filter sets for Hoechst and CellROX Deep Red signals, respectively. Only Calcein AM-positive live cells were analyzed for ROS signal to exclude oxidative stress artifacts from dead or dying cells.

**Cell Encapsulation in NorHA Hydrogels**

Hydrogels were cast as described previously. All hydrogel components were prepared under sterile conditions. Cells were encapsulated at a final density of $1 \times 10^7$ cells per mL hydrogel, and all hydrogels were cast at 25 µL volumes at 2 wt% with 35% norbornene modification. All cell handling and hydrogel fabrication procedures were performed in the dark under red light. Hydrogels were allowed to gel for 20 min at 37 °C to ensure complete network formation and cell encapsulation. Following confirmed gelation, hydrogels were submerged in DMEM and incubated at 37 °C, 5% $CO_2$.

**Live/dead Viability Assay of Encapsulated HDF (post softened) and NIH-3T3 cells**

Cell viability was assessed 24 hours post-encapsulation and/or light exposure using calcein acetoxymethyl ester (Calcein Green) and ethidium homodimer-1 (EtHD,). Hydrogels were washed 1x with PBS and submerged in PBS containing 2 µM Calcein and 4 µM EtHD. Hydrogels were incubated in staining solution at 37°C for 1 hour to allow adequate dye penetration throughout the hydrogel volume. Following incubation, hydrogels were washed twice with 1x PBS and imaged immediately. Calcein (live cells, green fluorescence) was imaged using the GFP filter set, and EtHD (dead cells, red fluorescence) was imaged using the mCherry filter set. Tiled scans were acquired across the hydrogel surface to ensure comprehensive sampling of the encapsulated cell population. Images were processed and analyzed using Imaris software. Cell viability was calculated as the percentage of calcein-positive cells relative to total cells (calcein-positive plus ethidium homodimer-1-positive). In the second experiment, the viability of encapsulated cells was assessed following photosoftening. Hydrogels were cast as described above, placed in the incubator for 30 min to ensure complete gelation, and then irradiated with 455 nm light (20 mW) for 10 min. Viability was subsequently measured by live/dead staining as described above. A minimum of three fields of view per hydrogel were analyzed. All experiments were performed with 3 biological and 3 technical replicates. Images were processed and analyzed using Imaris and ImageJ software **(Figure S18-S19).**

**CellROX Oxidative Stress Assay for Cell-Laden Hydrogels Before and After Photosoftening**

CellROX-based assay was used to assess oxidative stress in cells encapsulated within hydrogels under two conditions: in as-cast hydrogels and in Ru-HA hydrogels following photosoftening. For each condition, three formulation groups were prepared: negative control, positive control, and treatment. For the negative control, 100% photostable (diTet) hydrogels were prepared at 2 wt% HA-Nor (35% norbornene modification) and swelled in DMEM supplemented with 10 mM N-acetyl-L-cysteine (NAC). For the positive control, 100% photosensitive (RuTetrazine) hydrogels were prepared at 3 wt% HA-Nor (80% norbornene modification) to maximize ruthenium content and swelled in 10 mM sodium azide ($NaN_3$) in DMEM to induce oxidative stress. Treatment hydrogels were prepared at 2 wt% HA-Nor (35% norbornene modification) with 70:30 and 50:50 RuTetrazine:diTet crosslinker ratios, as these formulations demonstrated higher viability in prior experiments. To evaluate oxidative stress arising specifically from photosoftening, the RuTetrazine-containing treatment and positive-control hydrogels were allowed to gel for 30 min, overlaid with warm DMEM, and then immediately irradiated with 455 nm light (20 mW) for 10 min to photocleave RuTetrazine crosslinks and soften the network; the photostable NAC negative control was not irradiated, and the corresponding pre-softening hydrogels were handled identically but without irradiation. After casting (or irradiation, for the photosoftened groups), hydrogels were incubated at 37 °C for 24 h prior to staining.

A combined staining solution was prepared in warm DMEM containing CellROX Deep Red (5 µM final concentration, from 2.5 mM stock in DMSO; 2 µL per 1 mL media), Hoechst 33342 (5–10 µg $mL^{-1}$ final concentration, from 10 mg $mL^{-1}$ stock; 1 µL per 1 mL media), and calcein AM (2 µM final concentration, from 4 mM stock; 3 µL per 6 mL media). Media was aspirated from wells containing hydrogels and wells were washed once with PBS. Staining solution was added to fully cover each hydrogel (500 µL–1 mL depending on well size) and hydrogels were incubated at 37 °C for 1.5 h to allow dye diffusion through the macroporous network. Following incubation, the staining solution was aspirated and hydrogels were washed three times with PBS (10–15 min per wash) to allow excess dye to diffuse out. Fluorescence imaging was performed using DAPI (Hoechst 33342, blue nuclei) and Cy5 (CellROX Deep Red, ROS signal) filter cubes, and calcein AM signal (live cells) was imaged using a GFP/FITC filter cube. **(Figures S20-S21)**

**Confocal Imaging of Encapsulated cells in 80:20 RuTetrazine:diTet Hydrogels**

For the cell-spreading assay, HDFs or hMSCs were encapsulated at Day 0 in gels crosslinked with an 80:20 ratio of RuTetrazine:Ttz-MMP-RGDSK-Ttz to enable integrin-mediated adhesion and cell-directed matrix remodeling. When required for visualization of the polymer network, NorHA was prestained with Cy5-Tetrazine (Lumiprobe) prior to gelation. In brief: Cy5-tetrazine (2 µL of a 50 mM stock) was added to 30 mL of NorHA solution and allowed to react via iEDDA overnight, labeling a small fraction of the pendant norbornene groups. The labeled polymer was dialyzed against deionized water for 3 days to remove unreacted dye, then lyophilized and stored for use in hydrogel formulation.

Hydrogels were cast, incubated overnight, and photosoftened the following day (455 nm, 20 mW, 10 min). At the experimental endpoint (Day 10 for the spreading assay), hydrogels were washed with 1× PBS, fixed in 3.7% formaldehyde for 1 h at room temperature, and permeabilized in 0.5% Triton X-100 in PBS for 30 min. Gels were then blocked in blocking buffer. The blocking buffer was prepared by dissolving 90 mg BSA in 3 mL of 1× TBS containing 0.1% Tween 20 (3 µL Tween 20 per 3 mL), mixed gently until fully dissolved. Staining was performed in the same blocking buffer: Alexa Fluor 647-conjugated phalloidin (1:400) was used to visualize F-actin and Hoechst H3569 (1:1000) to label nuclei, applied together for 1 h at room temperature on the rocker. Diluting the dyes in blocking buffer rather than plain buffer suppresses background fluorescence and improves signal-to-noise in the macroporous network. Three 20-min PBS washes were performed between each step. Stained hydrogels were imaged on a Zeiss LSM 880 confocal microscope. For the spreading assay, Z-stacks spanning 500–1000 µm were acquired and processed in Fiji to generate maximum-intensity projections (MIPs) or rendered as 3D reconstructions in Imaris. MIPs were imported into Imaris, individual cells were segmented using the surface function, and cell surface area was extracted from the statistics module. Surface areas were compared between photosoftened and non-irradiated conditions in GraphPad Prism by unpaired Student's t-test.

**Figure S3. $^{1}$H NMR of Tet-COOH**

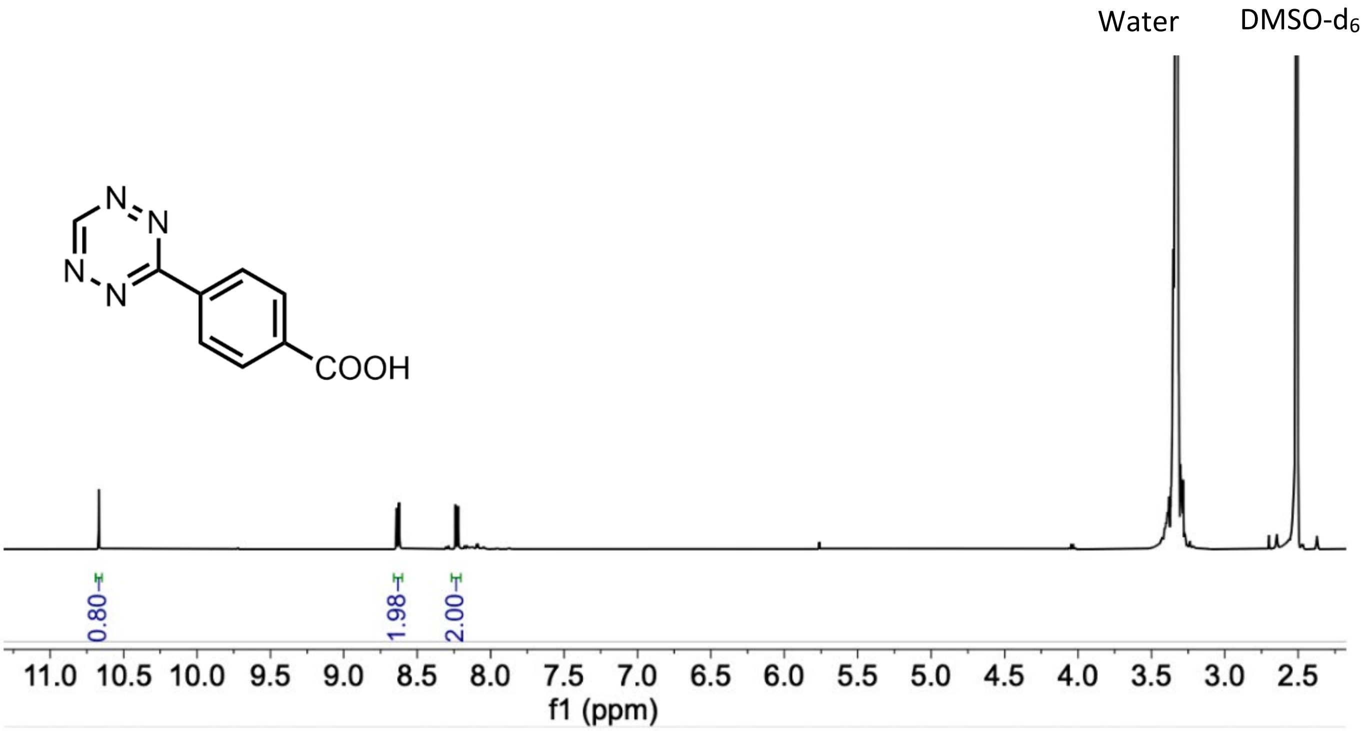

**Figure S3.** $^{1}$H NMR (DMSO-$d_6$) of tetrazine-COOH.

#### Figure S4. $^1$H NMR of TTZ-NHS

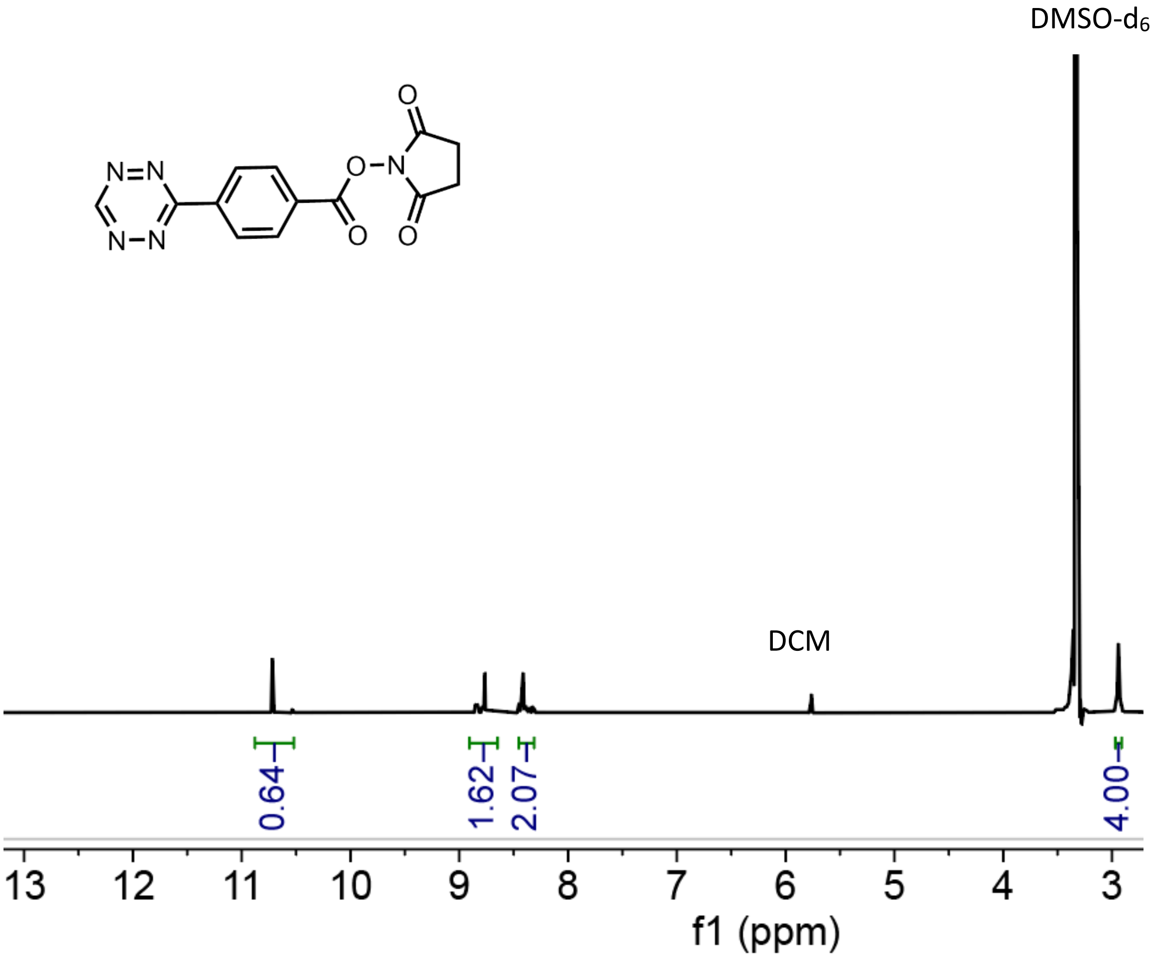

**Figure S4.** $^1$H NMR spectra (DSMO-d6) for TTZ-NHS ester

### Figure S5: 1H NMR of diTet

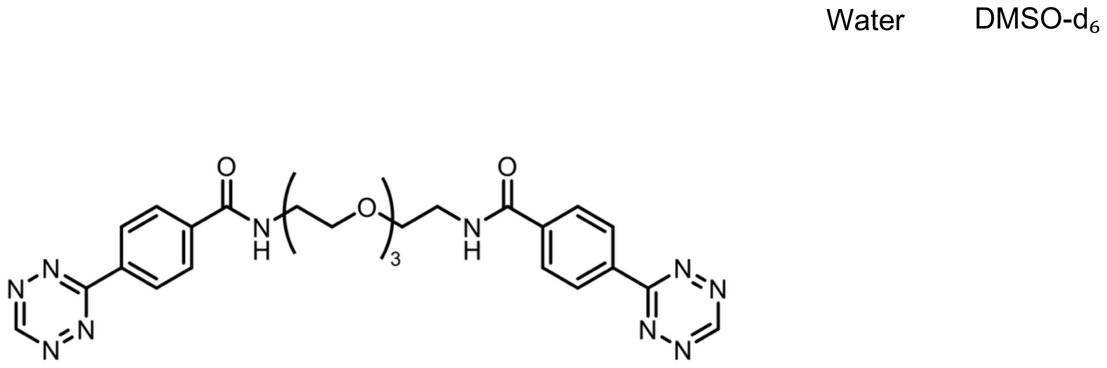

**Figure S5.** $^{1}H$ NMR spectra (DSMO-d6) for diTet

### Figure S6. 1H NMR of HA-nor

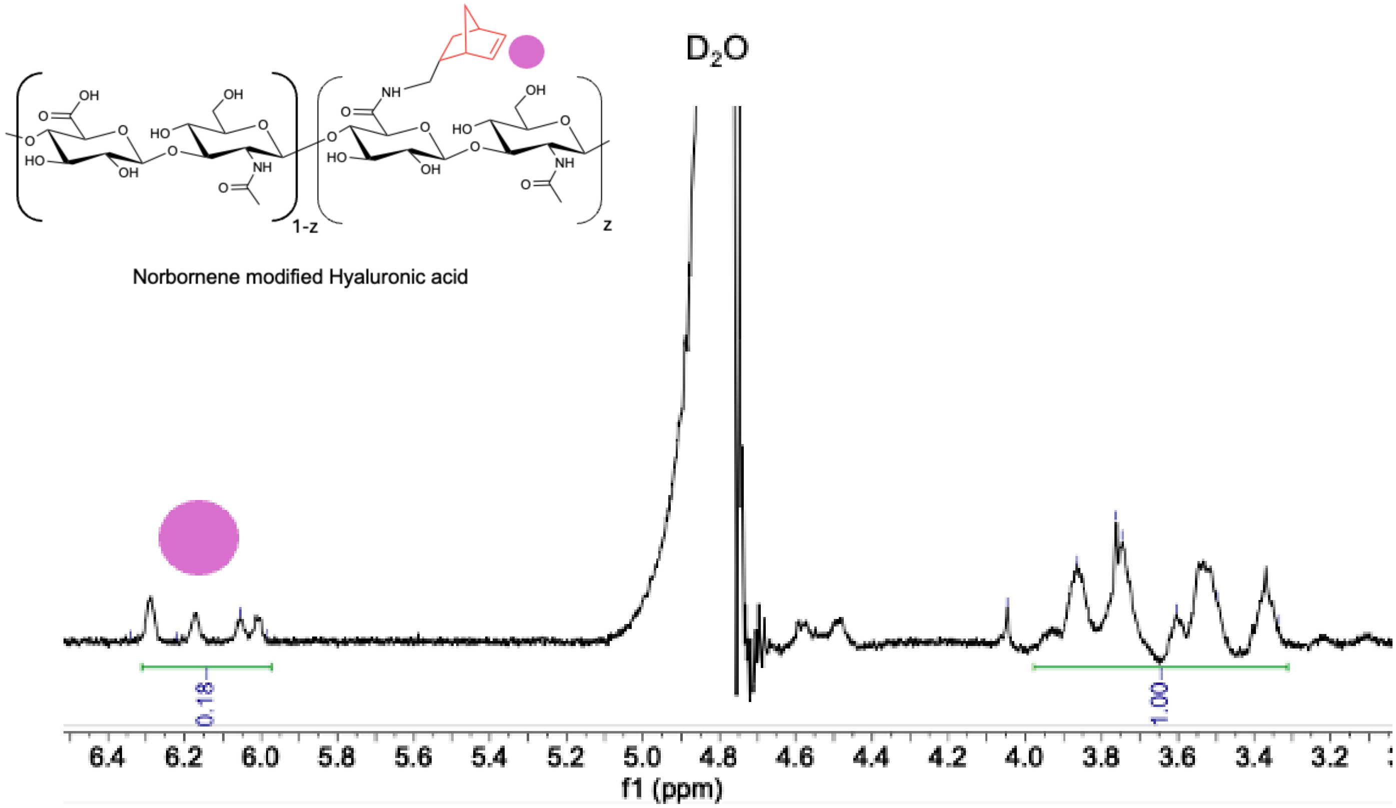

**Figure S6.** $^1$H NMR ($D_2O$) of NorHA to confirm modification and determine degree of substitution. NorHA was dissolved at approximately 7 mg/mL. The degree of substitution (%DS) was calculated by comparing the integration of the norbornene vinyl protons (δ = 5.90–6.30 ppm, 2H) relative to the HA backbone protons (δ = 3.20–4.10 ppm, 10H). In the above spectrum, the degree of substitution would be calculated as follows:$\left(\frac{0.18/2}{1/10}\right) * 100 = 90\%$

#### Figure S9. High-resolution ESI mass spectrometry of Ttz-MMP-RGDSK-Ttz

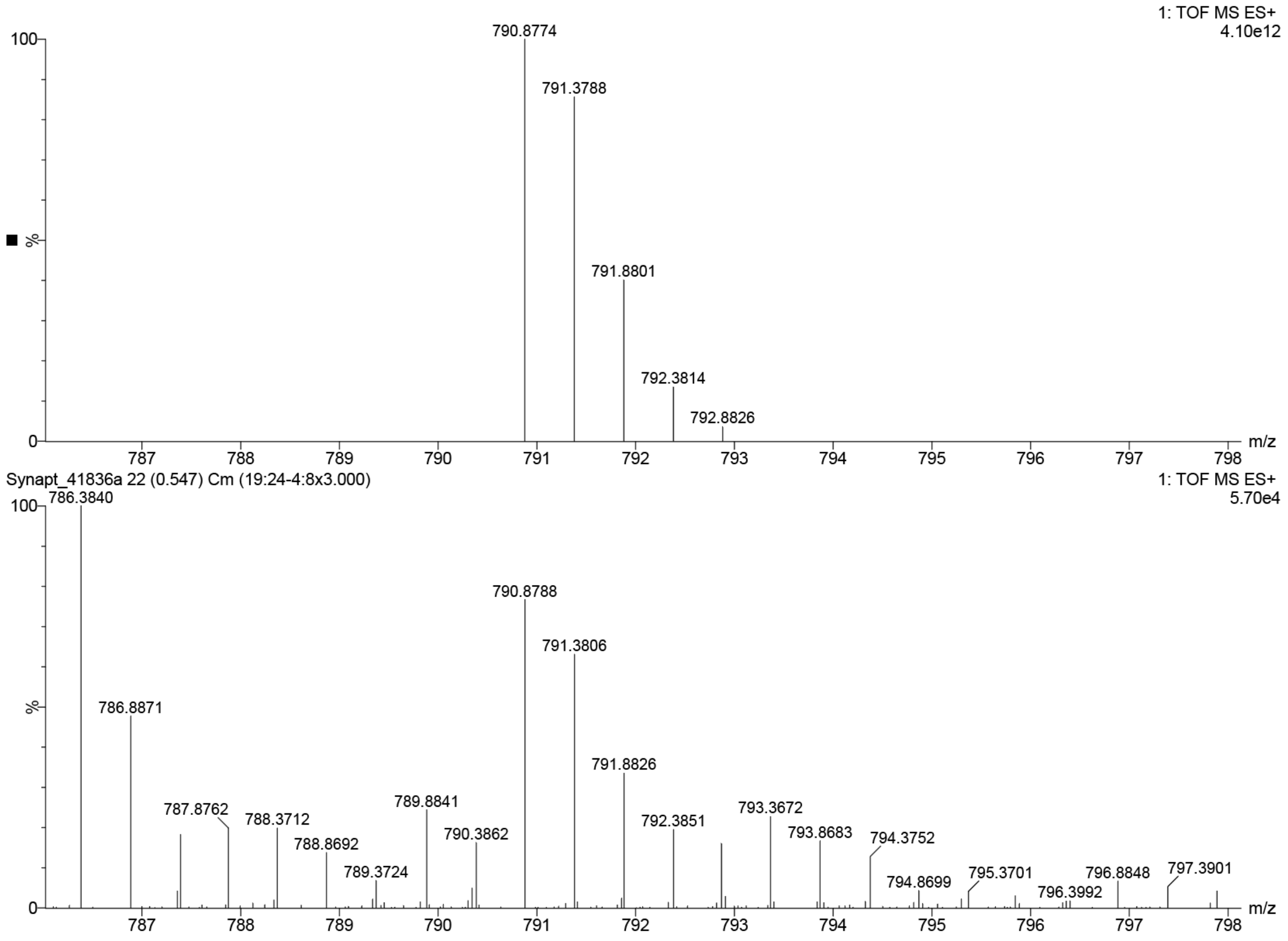

**Figure S9.** (Top) Theoretical isotope pattern calculated for $[C_{69}H_{99}N_{25}O_{19}]^{2+}$ (expected *m/2* = 790.8774; exact mass of neutral complex = 1579.74). (Bottom) Measured ESI-TOF mass spectrum of Ttz-MMP-RGDSK-Ttz (found *m/2* = 790.8788). The 0.5 *m/z* spacing of the isotope envelope confirms the 2+ charge state of the bis-tetrazine peptide, and agreement between theoretical and measured isotope distributions confirms the identity of the synthesized compound.

#### Figure S11. Photocleavage kinetics of RuTetrazine obey the Bunsen–Roscoe reciprocity law

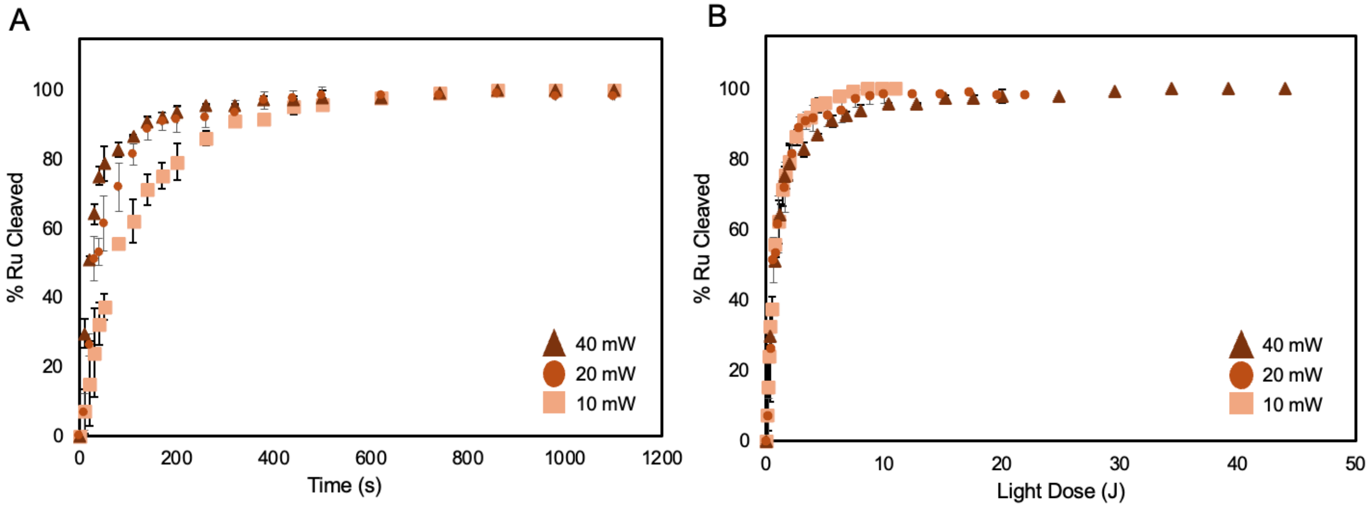

**Figure S11.** Photolysis of RuTetrazine (3 mM in 20:80 methanol:water) was monitored by UV-Vis absorbance spectroscopy under 450 nm irradiation at three power densities (10 mW, green triangles; 20 mW, orange squares; 40 mW, magenta circles). Percent RuTetrazine photocleaved was calculated from the decrease in the metal-to-ligand charge transfer (MLCT) absorbance band at 450 nm relative to the fully photolyzed endpoint. (A) Photocleavage as a function of irradiation time. Higher power densities yielded faster cleavage kinetics, with all conditions reaching ~100% cleavage within 600–1000 s. (B) The same data replotted against total light dose (fluence, J/cm², calculated as power × time / illuminated area) collapse onto a single master curve, reaching ~100% cleavage at approximately 2–3 J/cm² regardless of incident power. This convergence confirms that RuTetrazine photocleavage obeys the Bunsen–Roscoe reciprocity law, wherein photochemical outcome is governed by total photon dose rather than photon flux.

### Figure S12. IR of HA

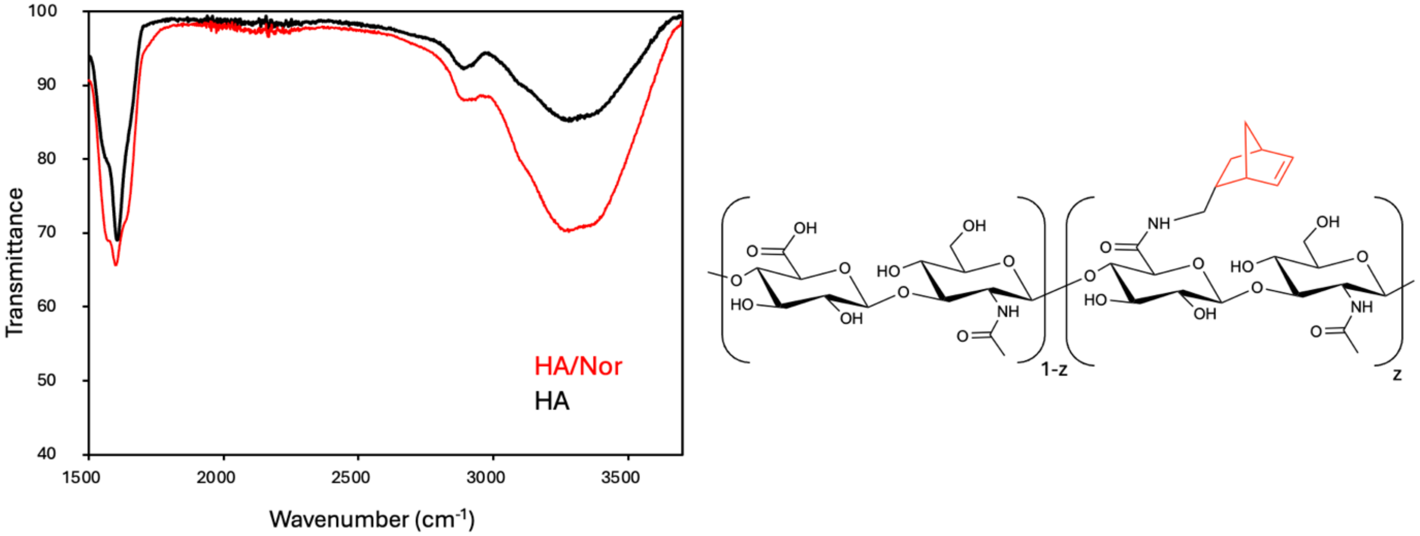

**Figure S12.** FTIR spectra were acquired using an ATR accessory (4000–400 $cm^{-1}$, 4 $cm^{-1}$ resolution). Successful norbornene conjugation was confirmed by increased N–H stretching absorption (3200–3400 $cm^{-1}$) in NorHA compared to unmodified HA, indicating amide bond formation.

#### Figure S13: Time sweep rheology of 3 wt% NorHA hydrogels

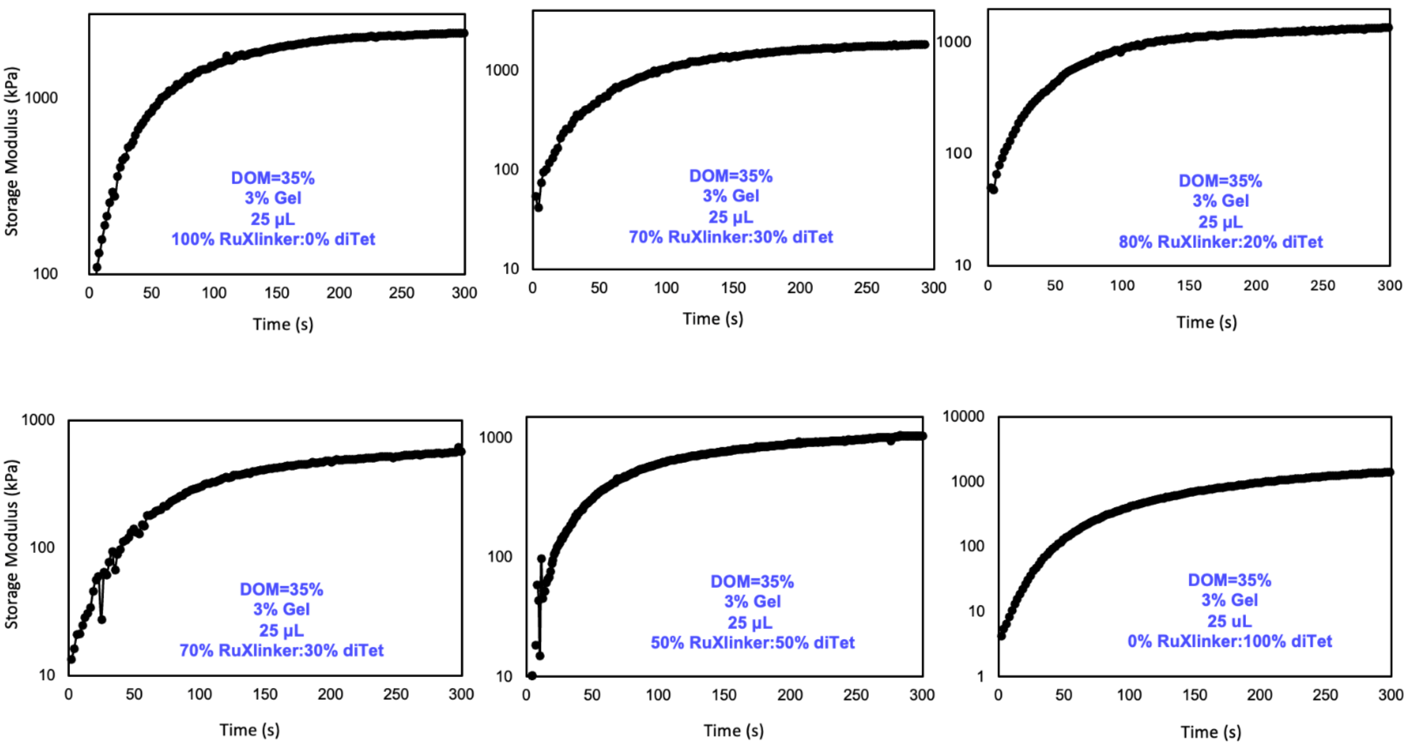

**Figure S13.** Time sweep rheology of 3 wt% NorHA hydrogels (35% norbornene modification, 25 µL) formed with varying RuTetrazine:DiTet crosslinker ratios (100:0, 80:20, 70:30, 50:50, and 0:100). Storage modulus (G′) was monitored during in situ gelation at 37 °C (1 Hz, 1% strain). All formulations reached their plateau modulus within 30 minutes, a gelation timescale compatible with cell encapsulation workflows.

#### Figure S14: Photosoftening 2wt% Hydrogels

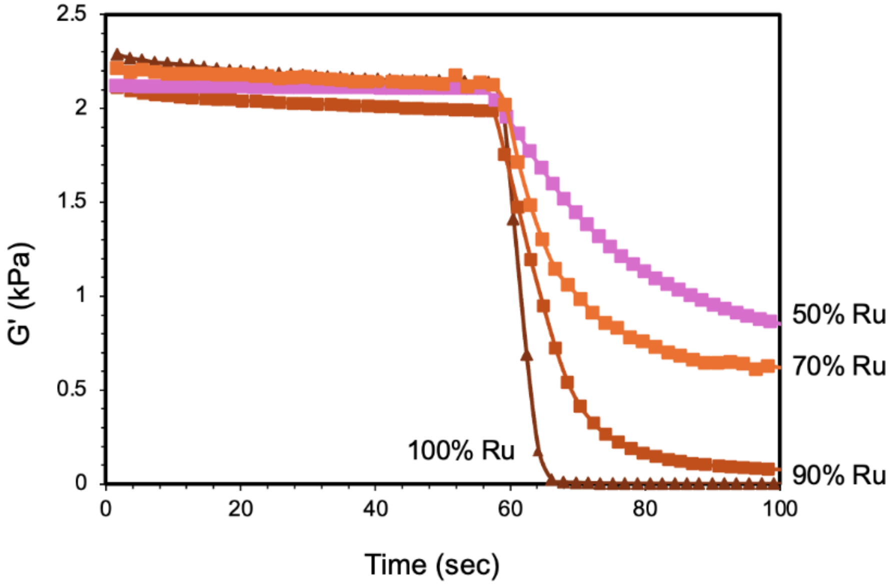

**Figure S14.** Photorheological characterization of 2 wt% Ru-HA hydrogels (35% norbornene modification) with varying RuTetrazine:DiTet crosslinker ratios (100:0, 90:10, 70:30, and 50:50). Storage modulus (G′) was monitored by oscillatory shear rheology (1 Hz, 1% strain) before, during, and after in situ irradiation with 455 nm light (20 mW). The 100% and 90% RuTetrazine formulations underwent near-complete softening, while formulations with higher DiTet content retained greater residual stiffness and required longer irradiation times, confirming tunable photosoftening through crosslinker composition.

**Figure S15: Photorheological characterization of an MMP-degradable, RGD-functionalized photosoftening hydrogel.**

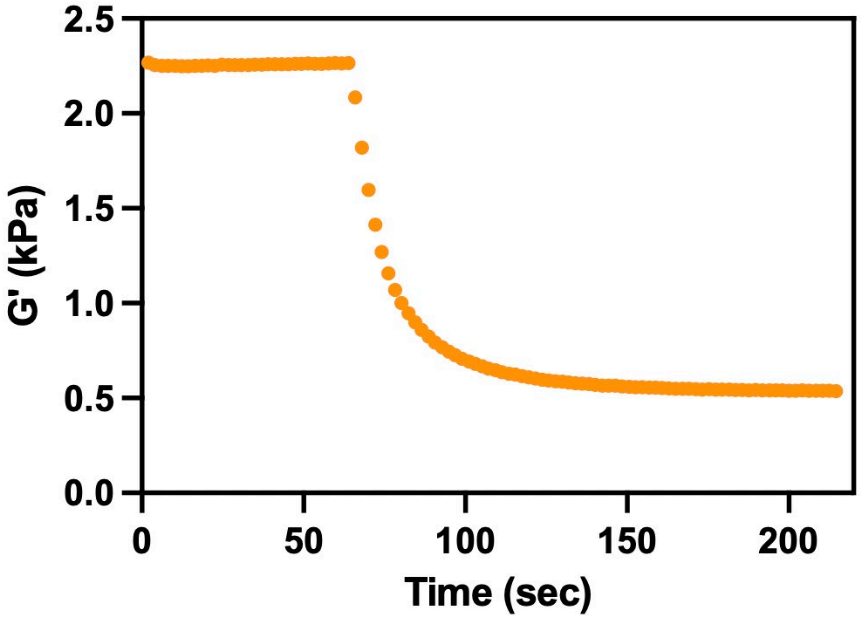

**Figure S15:** Storage modulus (G′) of a 2 wt% NorHA hydrogel (35% norbornene modification, 25 μL) crosslinked with an 80:20 ratio of RuTetrazine to Ttz-MMP-RGDSK-Ttz peptide, monitored by oscillatory shear rheology (1 Hz, 1% strain) at 37 °C. Following baseline equilibration, the gel was irradiated in situ with 455 nm light (20 mW, 5 min), beginning at ~60 s. G′ decreased from 2.27 kPa to 0.54 kPa. (Figure S17).

**Figure S16. IC50 of soluble RuTetrazine in HDFs.**

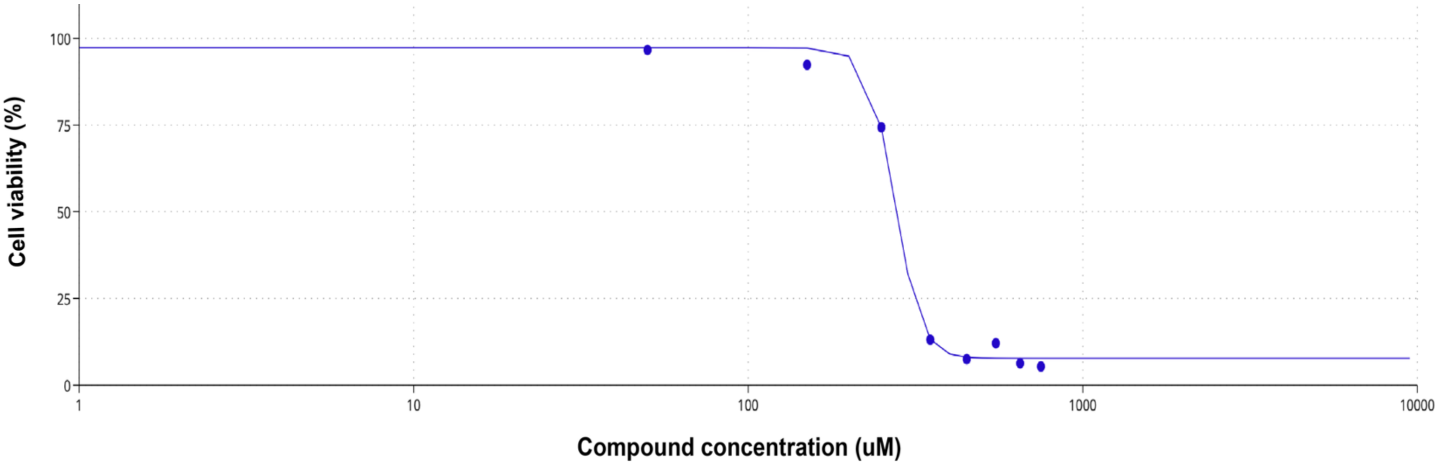

**Figure S16.** IC50 determination for soluble RuTetrazine in human dermal fibroblasts (HDFs). HDFs were exposed to RuTetrazine (0–750 µM) for 24 h, and metabolic activity was quantified by WST-8 assay and normalized to untreated controls. A four-parameter logistic fit (AAT Bioquest IC50 calculator) gave an IC50 of 274.8 µM.

**Figure S17. Ames mutagenicity assay of RuTetrazine**

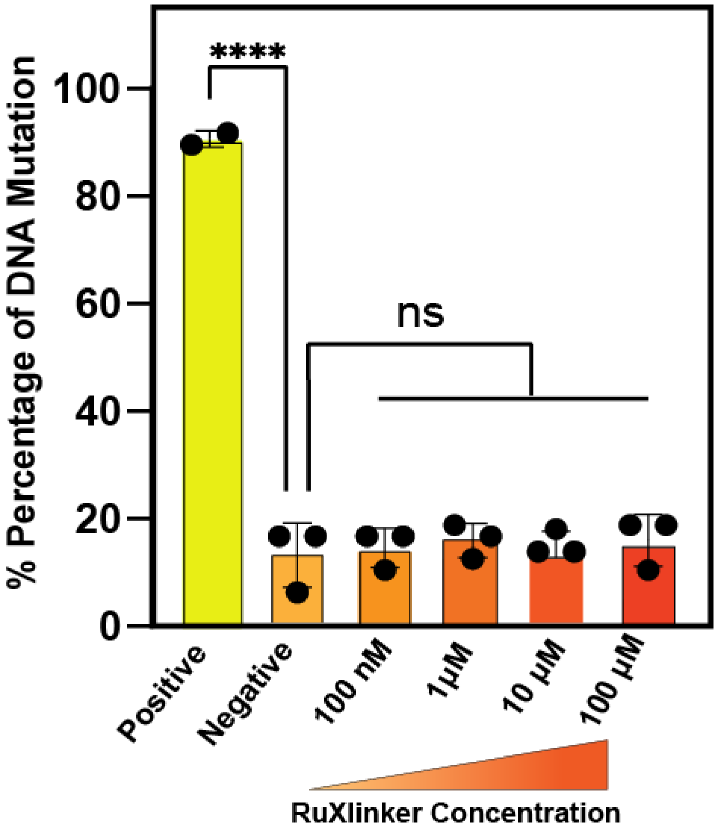

**Figure S17.** Ames mutagenicity assay of RuTetrazine across a concentration range. *Salmonella typhimurium* TA98 frameshift-reversion tester strain was exposed to RuTetrazine at 100 nM, 1 µM, 10 µM, and 100 µM [without S9 metabolic activation], and the percentage of DNA mutation was quantified from histidine-prototroph revertant colony formation. Bars from left: positive control (yellow), negative control (orange), and RuTetrazine at increasing concentrations (orange-to-red gradient). RuTetrazine showed no significant increase in mutation frequency over the negative control at any concentration tested. Data: mean ± SD; n = 3; ****$p < 0.0001$; ns = not significant (each RuTetrazine concentration versus negative control).

**Figure S18: Cytocompatibility of NIH-3T3 Fibroblasts Encapsulated in Ru-HA Hydrogels**

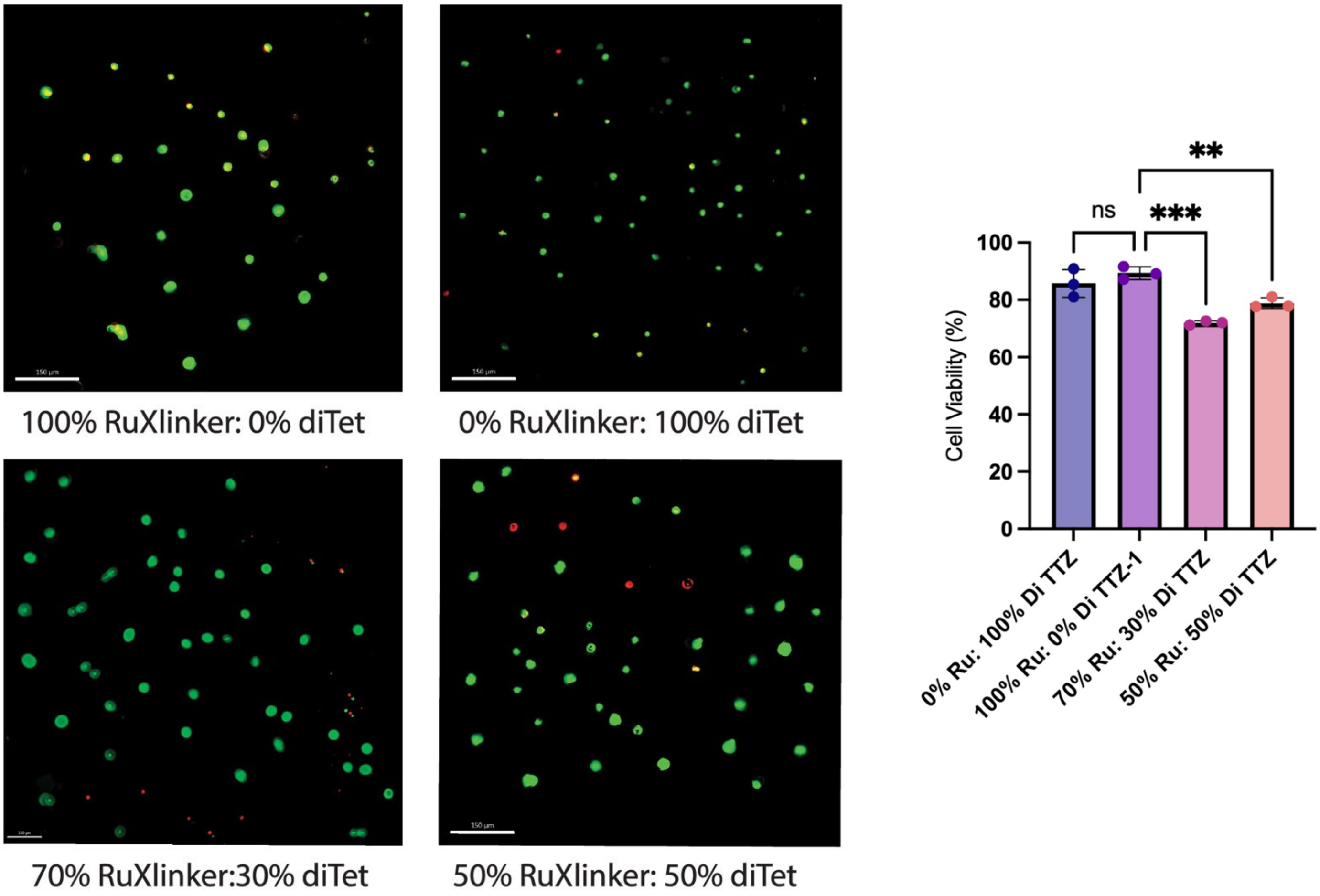

**Figure S18.** Live/dead viability of NIH-3T3 fibroblasts encapsulated in 3 wt% Ru-HA hydrogels with varying RuTetrazine:DiTet crosslinker ratios (90:10, 80:20, 70:30, and 50:50), a photostable control (0:100), and a fully photosensitive formulation (100:0), 24 hours post-encapsulation. Live cells (green, calcein AM) and dead cells (red, ethidium homodimer-1). All formulations exhibited >75% viability. Scale bars = 150 µm.

#### Figure S19: Cell Viability Following Photosoftening Hydrogels with Encapsulated NHDF

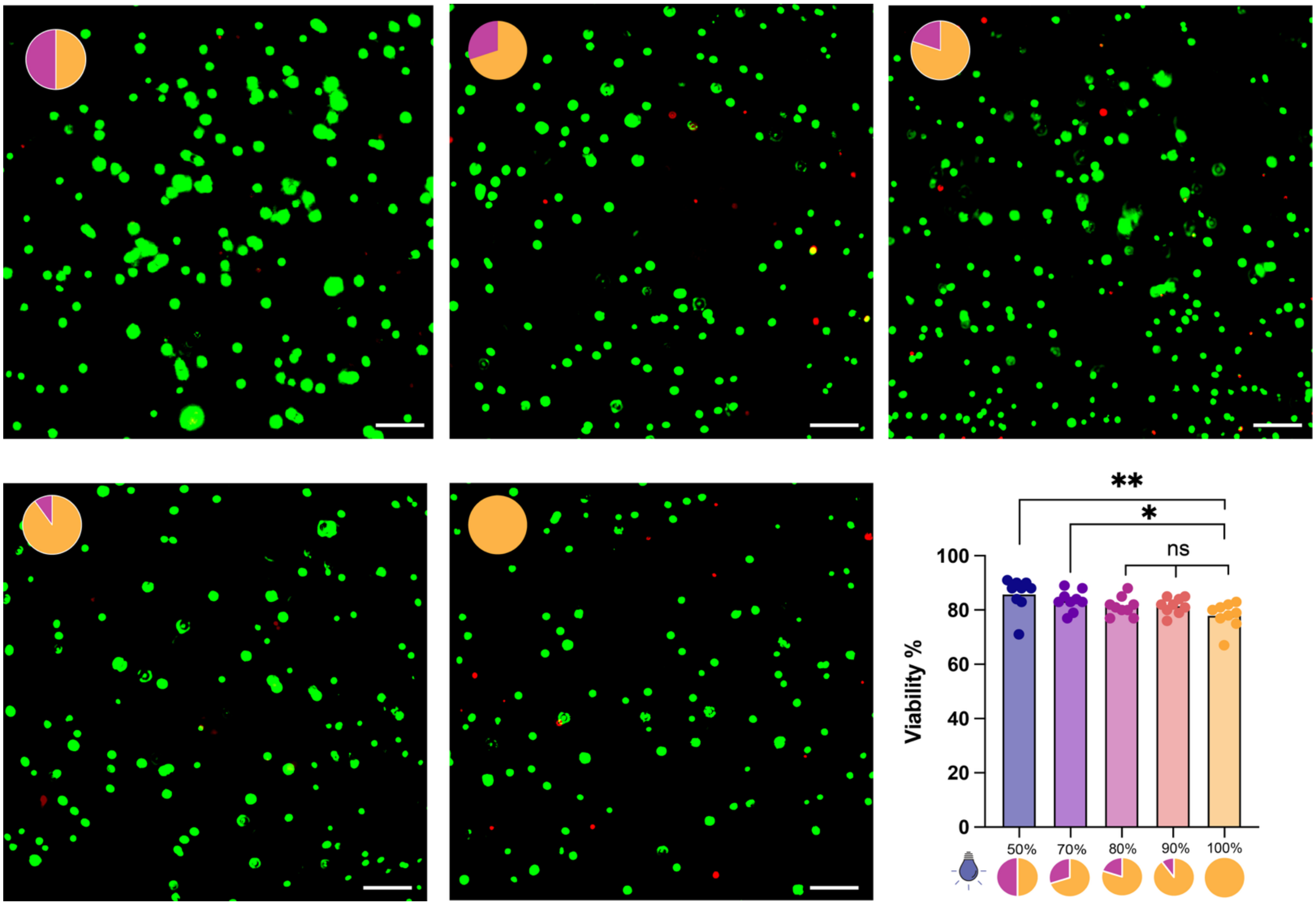

**Figure S19.** Live/dead viability of NHDFs encapsulated in 2 wt% Ru-HA hydrogels with varying RuTetrazine:DiTet crosslinker ratios (50:50, 70:30, 80:20, 90:10, and 100:0) 24 hours post-encapsulation. Live cells (green, calcein AM) and dead cells (red, ethidium homodimer-1) were imaged by widefield fluorescence microscopy. Pie charts indicate the proportion of photosensitive (orange) to photostable (pink) crosslinker in each formulation. All formulations exhibited >80% viability regardless of RuTetrazine content. Scale bars = 10

#### Figure S20: CellROX Oxidative Stress Assay in Hydrogels

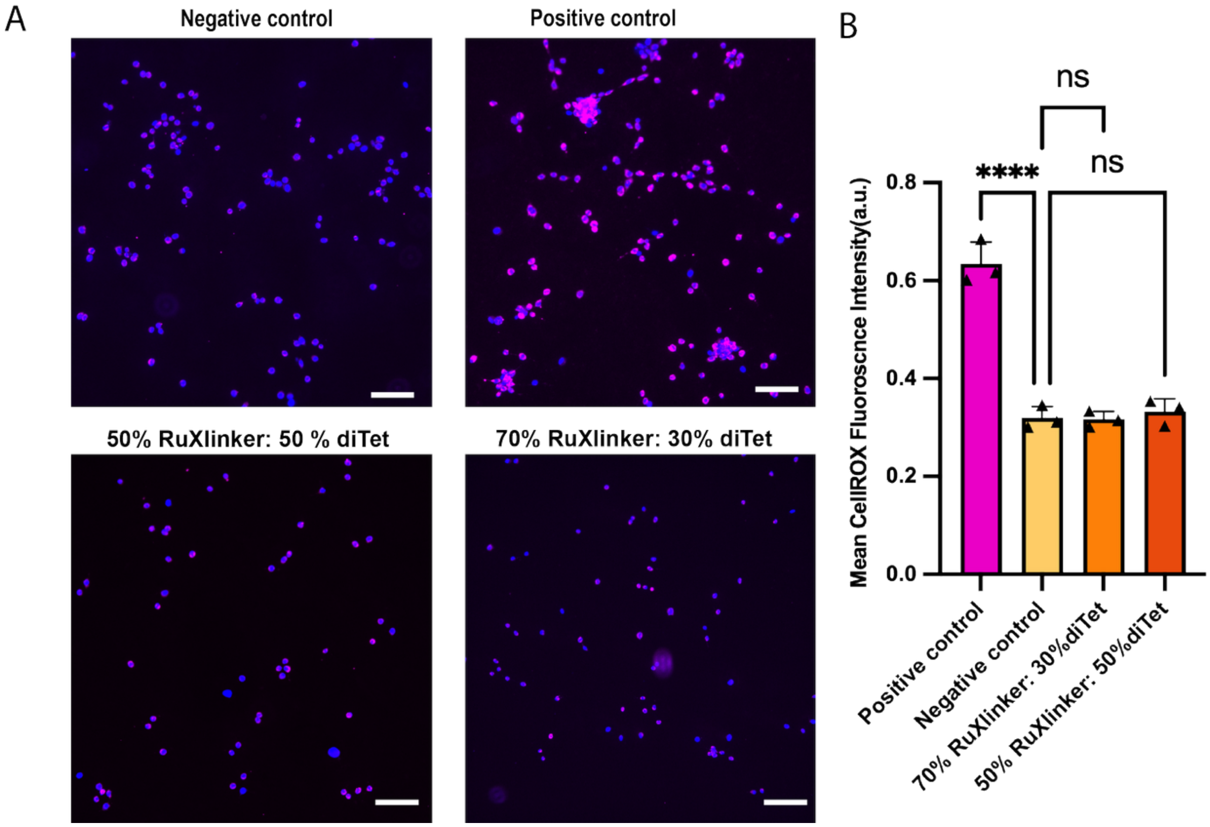

**Figure S20.** CellROX Deep Red oxidative stress assay of HDFs encapsulated in 2 wt% Ru-HA hydrogels (35% norbornene modification) with 70:30 and 50:50 RuTetrazine:DiTet crosslinker ratios. Negative control: 100% DiTet hydrogel (2 wt%) swelled in 10 mM *N*-acetyl-L-cysteine; positive control: 100% RuTetrazine hydrogel (3 wt%) swelled in 10 mM sodium azide. (A) Representative widefield fluorescence micrographs of encapsulated HDFs; nuclei marked by Hoechst 33342 (blue), intracellular ROS by CellROX Deep Red (magenta). Scale bars = 100 µm. (B) Mean CellROX fluorescence intensity per condition, quantified from calcein AM–positive live cells only. Data: mean ± SD; ***p < 0.001; ns = not significan

### Figure S21: CellROX Oxidative Stress Assay Following Photosoftening

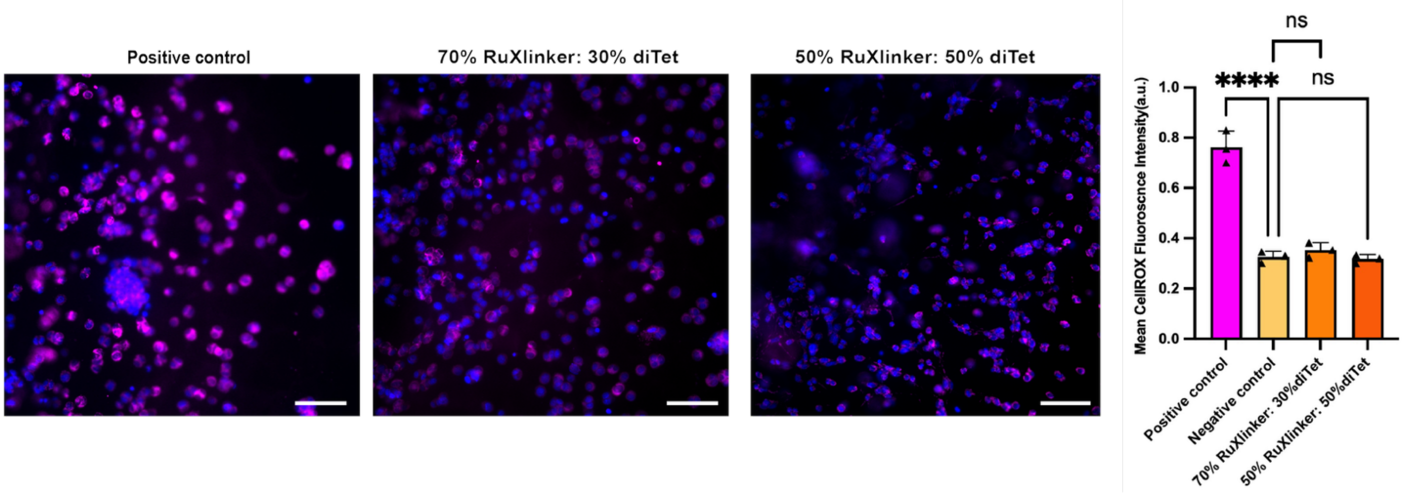

**Figure S21.** CellROX Deep Red oxidative stress assay of HDFs encapsulated in 2 wt% Ru-HA hydrogels (35% norbornene modification) with 70:30 and 50:50 RuTetrazine:diTet crosslinker ratios, assessed following photosoftening (455 nm, 20 mW, 10 min). Negative control: 100% DiTet hydrogel (2 wt%) swelled in 10 mM *N*-acetyl-L-cysteine; positive control: 100% RuTetrazine hydrogel (3 wt%) swelled in 10 mM sodium azide. (A) Representative widefield fluorescence micrographs of encapsulated HDFs; nuclei marked by Hoechst 33342 (blue), intracellular ROS by CellROX Deep Red (magenta). Scale bars = 100 µm. (B) Mean CellROX fluorescence intensity per condition, quantified from calcein AM–positive live cells only. Data: mean ± SD; **p $< 0.01$; ns = not significant.

**Figure S22. Confocal imaging of NHDFs at macropore boundaries in 3 wt% Ru-HA hydrogels.**

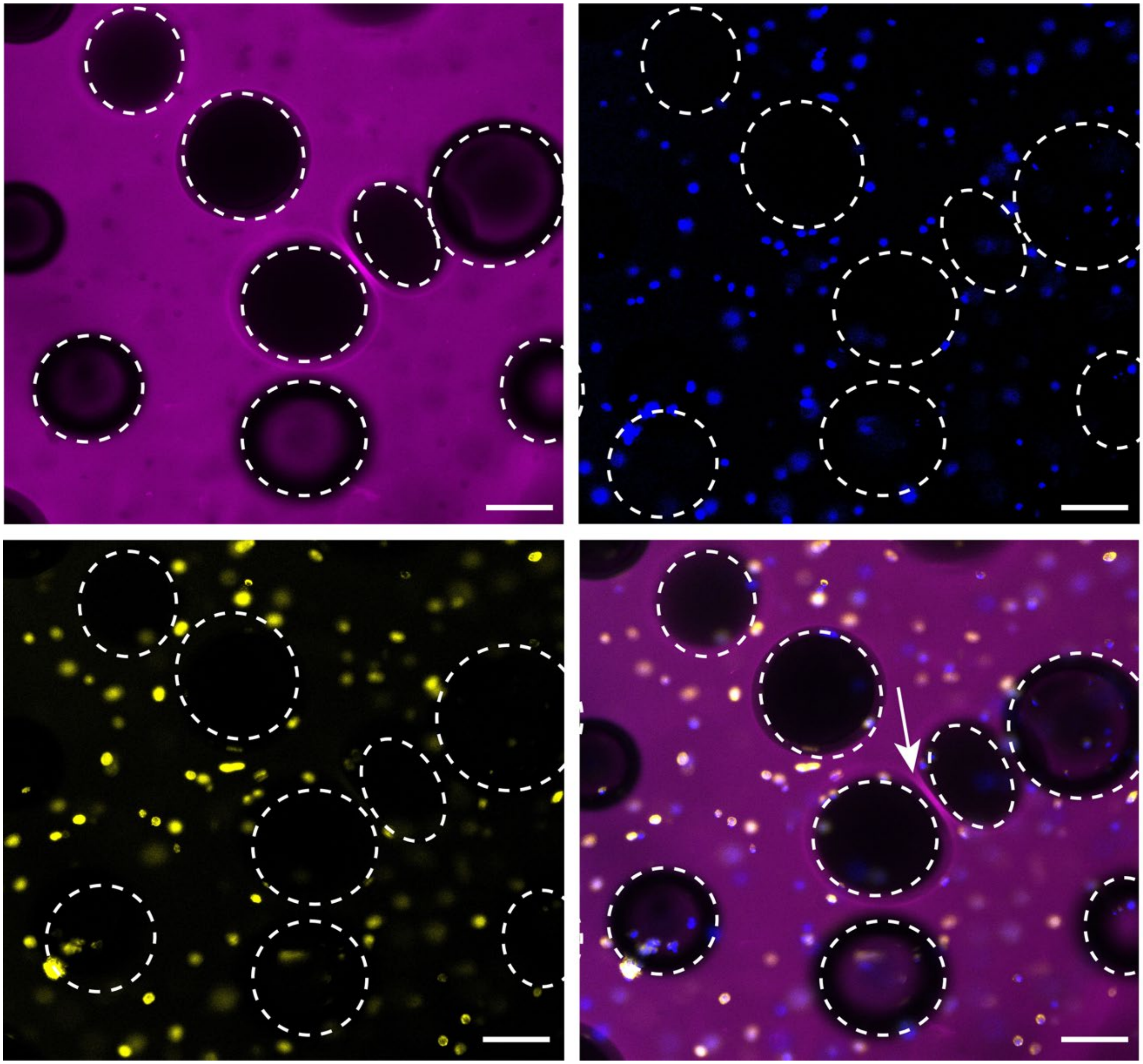

**Figure S22.** Additional confocal fluorescence micrographs of NHDFs encapsulated in 3 wt% Ru-HA hydrogels (35% norbornene modification, 100% RuTetrazine, imaged 24 hrs post-encapsulation), acquired on a Zeiss confocal microscope. Single channels show the polymer network labeled with Cy5-tetrazine (magenta, top left), nuclei stained with Hoechst (blue, top right), and F-actin labeled with phalloidin (yellow, bottom left); the merged image (bottom right). Dashed white circles outline representative macropores, which appear as dark circular exclusions across all channels. The white arrow indicates a region of densified polymer between two adjacent macropores, where the network was compacted as the gas bubbles expanded during gelation. Scale bars = 200 µm.

#### Figure S23: Bubble Size Quantification in Macroporous Ru-HA Hydrogels

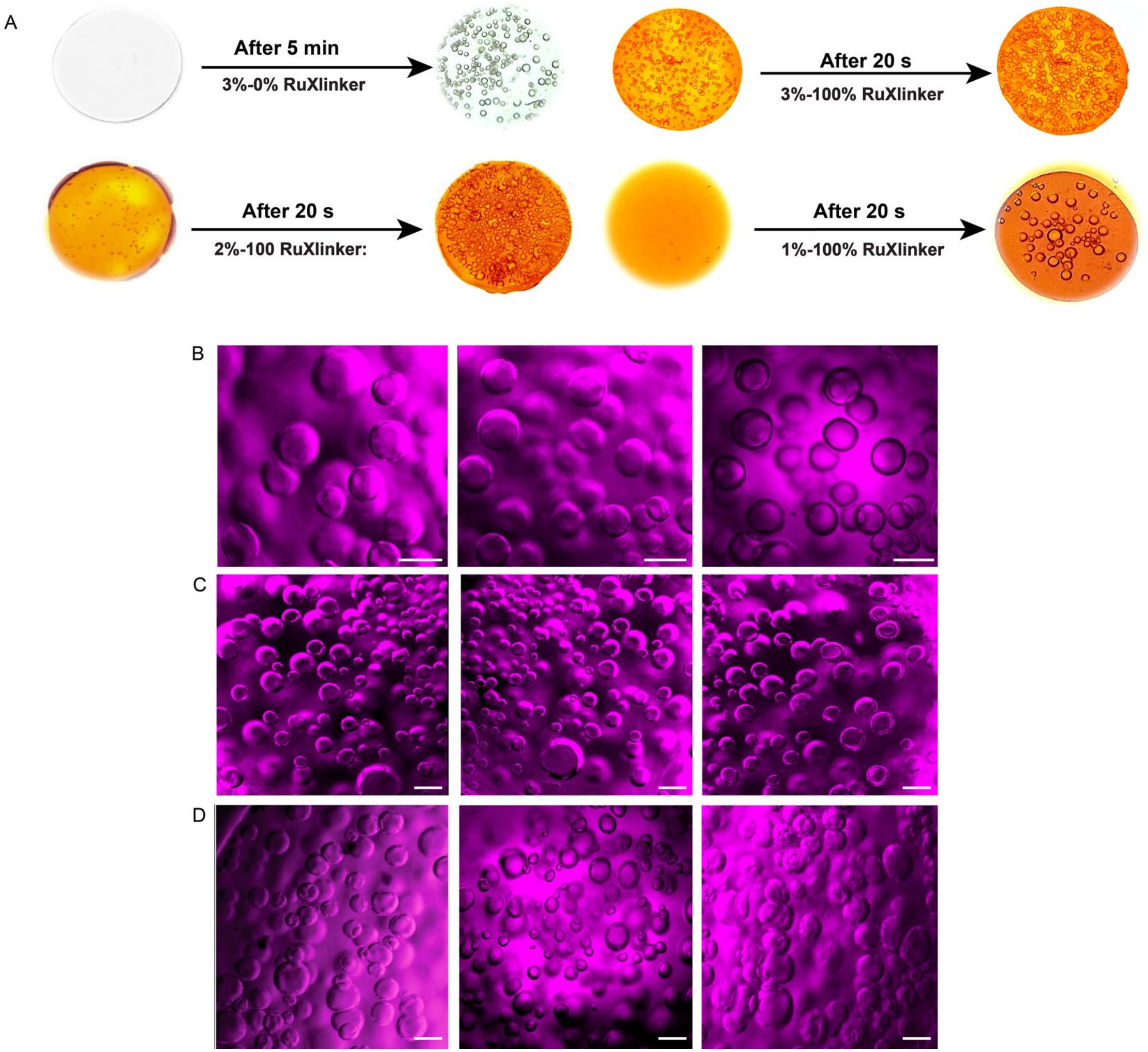

**Figure S23.** Macroporous structure of Ru-HA hydrogels generated by iEDDA-mediated $N_2$(g) evolution. A) Macroscopic images of hydrogels before and after gelation: 3 wt% with 100% DiTet (~20 min gelation), 3 wt% with 100% RuTetrazine (~5 min), 2 wt% with 100% RuTetrazine (~5 min), and 1 wt% with 100% RuTetrazine (~5 min). Bubble formation was observed in all formulations regardless of crosslinker identity. B–D) Fluorescence images of Cy5-tetrazine-labeled hydrogels at B) 3 wt%, C) 2 wt%, and D) 1 wt% HA-Nor (35% norbornene modification, 50 µL), with three representative fields of view per formulation. Scale bars = 100 µm.

#### Figure S24. Macropore stability over 10 days

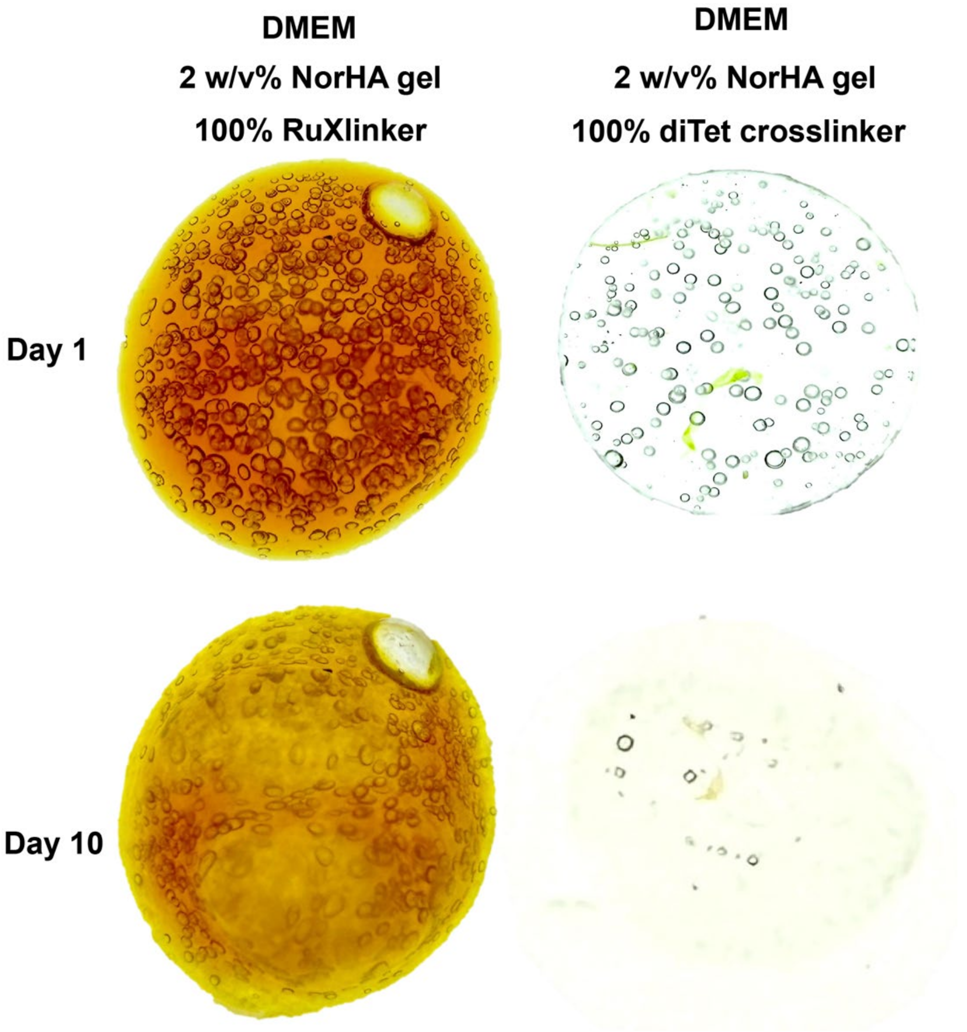

**Figure S24.** Macropore network under tissue-culture conditions. Stereomicroscope images of 2 w/v% NorHA hydrogels (35% norbornene modification) imaged at Day 1 and Day 10 of incubation at 37 °C. Left: 100% RuTetrazine hydrogel in DMEM. Right: 100% diTet hydrogel in PBS.
